# BDSF is a degradation-prone quorum-sensing signal detected by the histidine kinase RpfC of *Xanthomonas campestris* pv. *campestris*

**DOI:** 10.1101/2022.01.03.474871

**Authors:** Xiu-Qi Tian, Yao Wu, Zhen Cai, Wei Qian

**Affiliations:** State Key Laboratory of Plant Genomics, Institute of Microbiology, Chinese Academy of Sciences, Beijing 100101, China; School of Life Sciences, University of Chinese Academy of Sciences, Beijing 100049, China; Aviation General Hospital of China Medical University & Beijing Institute of Translational Medicine, Chinese Academy of Sciences, Beijing 100012, China; College of Advanced Agricultural Sciences, University of Chinese Academy of Sciences, Beijing 100049, China; CAS Center for Excellence in Biotic Interactions, University of Chinese Academy of Sciences, Beijing 100049, China

**Keywords:** Quorum-sensing, DSF, BDSF, *Xanthomonas campestris*, histidine kinase, functional divergence

## Abstract

Diffusible signal factors (DSFs) are medium-chain fatty acids that induce bacterial quorum sensing. Among these compounds, BDSF is a structural analog of DSF that is commonly detected in bacterial species (e.g., *Xanthomonas*, *Pseudomonas*, and *Burkholderia*). Additionally, BDSF contributes to the interkingdom communication regulating fungal life stage transitions. How BDSF is sensed in *Xanthomonas* spp. and the functional diversity between BDSF and DSF remain unclear. In this study, we generated genetic and biochemical evidence that BDSF is a low-active regulator of *X. campestris* pv. *campestris* quorum sensing, whereas *trans*-BDSF seems not a signaling compound. BDSF is detected by the sensor histidine kinase RpfC. Although BDSF has relatively low physiological activities, it binds to the RpfC sensor with a high affinity and activates RpfC autophosphorylation to a level that is similar to that induced by DSF *in vitro*. The inconsistency in the physiological and biochemical activities of BDSF is not due to RpfC– RpfG phosphorylation or RpfG hydrolase. Neither BDSF nor DSF controls the phosphotransferase and phosphatase activities of RpfC or the ability of RpfG hydrolase to degrade the bacterial second messenger cyclic di-GMP. We demonstrated that BDSF is prone to degradation by RpfB, a critical fatty acyl-CoA ligase involved in the turnover of DSF-family signals. *rpfB* mutations lead to substantial increases in BDSF-induced quorum sensing. Although DSF and BDSF are similarly detected by RpfC, our data suggest that their differential degradation in cells is the major factor responsible for the diversity in their physiological effects.

**IMPORTANCE:** Diffusible signal factor (DSF) family are quorum-sensing signals employed by gram-negative bacteria. These signals are a group of *cis*-2-unsaturated fatty acids, such as DSF, BDSF, IDSF, CDSF, and SDSF. However, the functional divergence of various DSF signals remains unclear. The present study demonstrates that though BDSF is a low active quorum-sensing signal than that of DSF, it binds histidine kinase RpfC with a higher affinity and activates RpfC autophosphorylation to the similar level as DSF. Rather than regulation of enzymatic activities of RpfC and its cognate response regulator RpfG encoding a c-di-GMP hydrolase, BDSF is prone to degradation in bacterial cells by RpfB, which effectively avoided the inhibition of bacterial growth by accumulating high concentration of BDSF. Therefore, our study shed new light on the functional difference of quorum-sensing signals and revealed that bacteria balance quorum-sensing and growth by fine-turning concentration of signaling chemical.

## INTRODUCTION

Diffusible signal factors (DSFs) are important quorum-sensing signals produced and detected by many Gram-negative bacteria (1, 2). The DSF-family signals form a group of medium-chain fatty acids, including DSF (*cis*-11-methyl-dodecenoic acid), BDSF (*cis*-2-dodecenoic acid), CDSF (*cis*,*cis*-11-methyldodeca-2,5-dienoic acid), IDSF (*cis*-10-methyl-2-dodecenoic acid), and SDSF (*trans*-2-decenoic acid) (3–5). Among these signaling molecules, DSF was the first to be identified in the phytopathogenic bacterium *Xanthomonas campestris* pv. *campestris*. In most *Xanthomonas* species, a conserved *rpf* gene cluster (encoding RpfC, RpfG, RpfB, and RpfF) is tightly associated with DSF-induced quorum sensing (1, 5, 6). The histidine kinase RpfC and the response regulator RpfG form a two-component signal transduction system. As a quorum-sensing signal, DSF binds to the N-terminal sensor region of RpfC to release the autoinhibition of autokinase activity (7). Next, RpfC putatively phosphorylates the response regulator RpfG, which has a C-terminal HD-GYP domain that acts as a hydrolase to degrade the second messenger Bis-(3′-5′)-cyclic dimeric guanosine monophosphate (c-di-GMP) into GMP (8). On the basis of protein phosphorylation, the RpfC–RpfG system positively regulates the production of various virulence factors, including extracellular polysaccharides (EPS) and extracellular enzymes, but negatively controls the production of DSF (9, 10). A recent study indicated RpfF is a key enzyme for synthesizing DSF because of its 3-hydroxyacyl-acyl carrier protein dehydratase and thioesterase activities. A mutation in *rpfF* eliminates the ability of bacteria to synthesize DSF-family signals (11). The genetic inactivation of *rpfC*, *rpfG*, and *rpfF* significantly decreases the virulence and biofilm development of *X. campestris* pv. *campestris* (5, 12). Additionally, the fatty acyl-CoA ligase activity of RpfB contributes to the degradation and turnover of DSF-family signals. Unlike other *rpf* genes, the inactivation of *rpfB* slightly increases the virulence of phytopathogenic bacteria (13, 14).

In contrast to the DSF chemical structure, BDSF (*cis*-dodecenoic acid) lacks a methyl group at the C-11 site and it has an isomeric conformation (*cis* and *trans* forms). Additionally, BDSF, which was first identified in the Gram-negative bacterium *Burkholderia cenocepacia*, helps regulate bacterial biofilm formation, motility, and virulence (4). Interestingly, both BDSF and *trans*-BDSF modulate germ tube formation in the fungus *Candida albicans*, implying BDSF can mediate interkingdom communication (15, 16). Both DSF and BDSF are commonly produced by *Xanthomonas* species. During an infection of cabbage, *X. campestris* pv. *campestris* uses diverse plant-derived compounds to synthesize more BDSF than DSF. Accordingly, BDSF may be a major *in planta* quorum-sensing signal during bacterium–plant interactions (17). In *B. cenocepacia*, the histidine kinase BCAM0227 is a potential BDSF receptor; the inactivation of its coding sequence partially disrupts BDSF-induced regulation (18). There is no biochemical evidence of the BDSF–BCAM0227 interaction. Moreover, a *B. cenocepacia* RpfR, which contains PAS, GGDEF, and EAL domains, was identified as a BDSF receptor [dissociation constant (*K*_d_) of 8.77 × 10^−7^ M] (19, 20). A structural analysis revealed that BDSF binds to the RpfR PAS domain at Asn^202^ (21). The RpfR GGDEF and EAL domains participate in the turnover of the bacterial second messenger c-di-GMP. Furthermore, BDSF and c-di-GMP control the formation of the RpfR–GtrR complex to modulate the GtrR transcription factor activity (22).

*Xanthomonas campestris* pv. *campestris* has genes encoding eight proteins with PAS, GGDEF, and EAL domains, but it lacks a full-length RpfR ortholog and its BDSF receptor remains unknown. Because RpfC is a DSF receptor and there is only a slight structural difference between DSF and BDSF, RpfC is most likely the BDSF receptor in *X. campestris* pv. *campestris*, but this will need to be experimentally verified. Even if RpfC is the BDSF receptor, the biochemical difference between DSF–RpfC and BDSF–RpfC interactions and their biological effects will need to be determined. In the present study, we demonstrated that BDSF, rather than *trans*-BDSF, is biologically active in *X. campestris* pv. *campestris* and that BDSF binds to the N-terminal region of RpfC. Both DSF and BDSF similarly activate the RpfC autokinase, but BDSF is substantially less biologically active than DSF. This difference is unrelated to the RpfC autokinase, phosphotransferase, and phosphatase activities induced by DSF or BDSF. We demonstrated that the *in vivo* degradation of BDSF by RpfB is the major reason for the difference between the two signals, which alleviated the inhibition of bacterial growth by high concentration of BDSF.

## RESULTS

### BDSF is less biologically active than DSF

Although BDSF and DSF are structural analogs, BDSF does not have a methyl group covalently attached to the C-11 site (Fig. 1A). Of the two BDSF isomers, the *cis* form is primarily produced by bacteria under natural conditions (23). To investigate the biological differences between BDSF and DSF in terms of their ability to induce quorum sensing, an *rpfF* null mutant (Δ*rpfF*), which cannot synthesize DSF-family molecules, was used to examine quorum sensing-related phenotypic changes. Without DSF and BDSF, the *rpfF* mutant did not produce and secrete extracellular proteases (Fig. 1B, left colony in each panel) and only limited biofilm formation was detected by crystal violet staining (Fig. 1C and 1D). The addition of DSF at low concentrations (0.5–5 µM) significantly increased the production of extracellular proteases and biofilm development, which was consistent with the results of previous studies (Fig. 1B and 1C) (3, 7). However, to induce similar phenotypic changes, BDSF required a higher concentration than DSF. More specifically, for the extracellular protease activity assay, the effect of 100–200 µM BDSF was equivalent to that of 5 µM DSF (Fig. 1B). For the biofilm quantification assay, the biofilm production induced by 0.5 µM DSF was roughly equal to the inductive effect of >10 µM BDSF (Fig. 1C and 1D).

**FIG 1.**
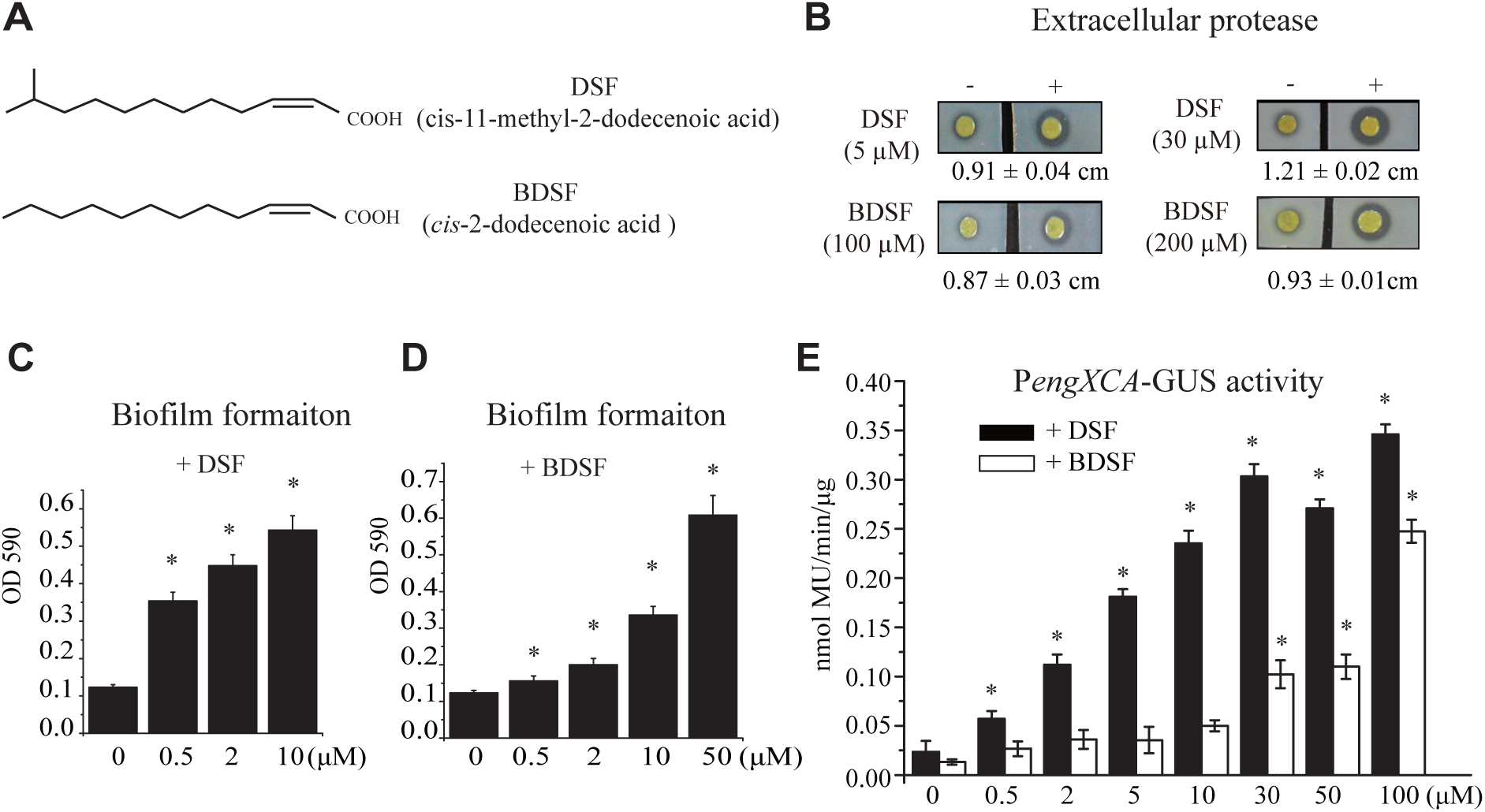
The quorum-sensing activity of BDSF is lower than that of DSF. (A) Chemical structures of BDSF and DSF. (B) Addition of BDSF and DSF increased the extracellular protease production of the *rpfF* mutant. In each panel, *rpfF* mutant bacterial colonies on NYG medium are presented, with the effects of 4 μL BDSF or DSF added nearby the *rpfF* mutant to trigger quorum sensing presented on the right. Protease activity was determined by measuring the diameters (cm) of the protein degradation zones after a 36-h incubation (n = 4). (C and D) Addition of DSF (C) and BDSF (D) increased the biofilm formation of the *rpfF* mutant. A crystal violet staining method was used to quantify the biofilm formation (n = 4). (E) *engXCA* transcription in response to BDSF or DSF. Each experiment was repeated four times. The vertical bar represents the standard deviation (n = 4). The asterisk indicates a significant difference compared with the unstimulated control.

In *X. campestris* pv. *campestris*, the expression of a gene encoding the extracellular endoglucanase EngXCA is regulated by RpfC–RpfG and DSF signals. The *engXCA* transcript level has traditionally been used for quantifying DSF activity (24, 25). In the current study, we used the *rpfF* mutant containing a reporter construct comprising the *engXCA* promoter fused to a β-glucuronidase (*gus*) gene (P*engXCA*-*gus*) to quantify the DSF and BDSF biological activities. The addition of various concentrations of DSF (0.5–100 µM) significantly increased the P*engXCA*-*gus* activity, with 0.5 µM DSF sufficient for inducing a 2.4 times increase in activity (relative to the unstimulated control) (Fig. 1E). However, the addition of BDSF (0.5 µM to 10 µM) only slightly affected P*engXCA*-*gus* activity. Similar to the biofilm assay, the biological activity of 100 µM BDSF was equivalent to that of 10 µM DSF. Considered together, these results revealed that BDSF is a low-activity signal that triggers *X. campestris* pv. *campestris* quorum sensing.

### *trans*-BDSF is an exceptionally low-activity regulator of bacterial quorum sensing

There are two isomers of the BDSF signal (i.e., *cis* and *trans*). Previous research indicated that both isomers can regulate germ tube and biofilm development in *C. albicans* (16); however, the biological activities of these isomers were not characterized in *X. campestris* pv. *campestris*. The aforementioned assays were conducted to compare the regulatory effects of BDSF and *trans*-BDSF on quorum sensing. The addition of high concentrations (50–200 µM) of *trans*-BDSF did not substantially affect the production of extracellular proteases or biofilm development (Fig. 2A and 2B). For the P*engXCA*-*gus* reporter assay, the addition of 50 and 100 µM *trans*-BDSF increased the P*engXCA*-*gus* activity level to 1.4- and 2.8 times that of the unstimulated control, respectively, which was significantly lower than the activity induced by BDSF (8.4 and 18.9 times, respectively, Fig. 2C). Therefore, the regulatory effects of *trans*-BDSF on *X. campestris* pv. *campestris* quorum sensing are much lower than those of BDSF.

**FIG 2.**
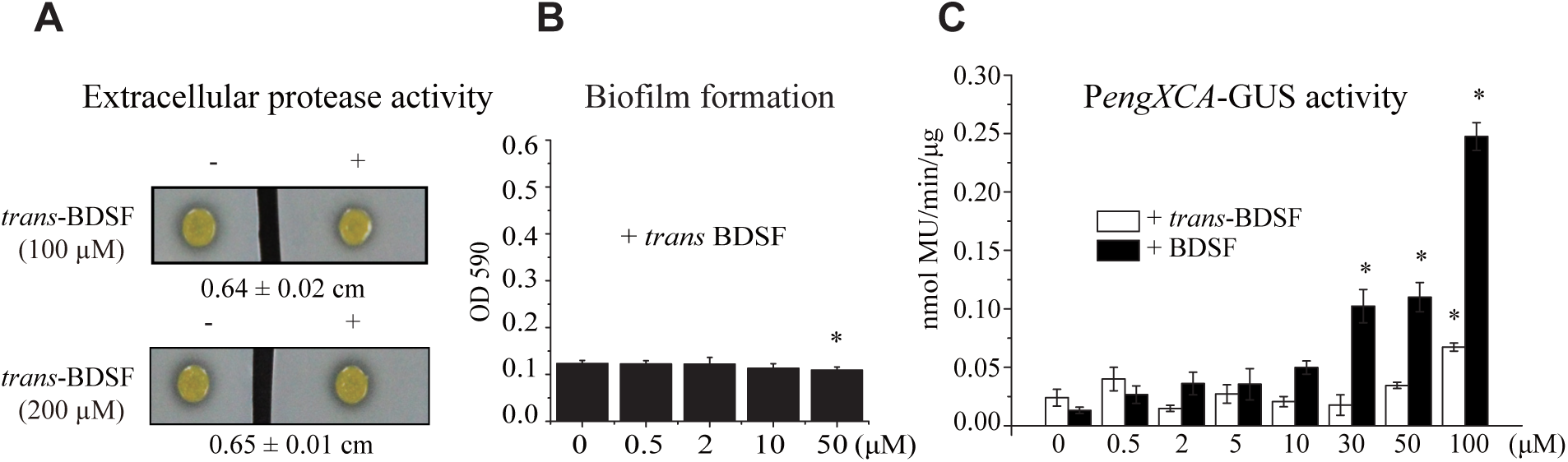
*trans*-BDSF is a low-activity inducer of quorum sensing. (A) Extracellular protease production assay. In each panel, *rpfF* mutant bacterial colonies on NYG medium are presented, with the effects of 4 μL *trans*-BDSF added nearby the *rpfF* mutant to trigger quorum sensing presented on the right. Protease activity was determined by measuring the diameters (cm) of the protein degradation zones after a 36-h incubation (n = 4). (B) Biofilm production. A crystal violet staining method was used to quantify the biofilm formation of the *rpfF* mutant (n = 4). (C) P*engXCA* reporter activity assay in response to BDSF or *trans*-BDSF. Each experiment was repeated four times. The vertical bar represents the standard deviation (n = 4). The asterisk indicates a significant difference compared with the unstimulated control.

### RpfC sensor and transmembrane regions are critical for sensing BDSF

RpfC is a hybrid histidine kinase with a short predicted sensor comprising 22 amino acid residues and five transmembrane (TM) helices at the N-terminal (Fig. 3A). We previously revealed that the signal input region (including the sensor and TM) is critical for detecting DSF (7). In this study, we employed multiple domain-deletion mutants and 21 mutants with point mutations (all in the Δ*rpfF* background) that were created via alanine-scanning mutagenesis to identify the protein region and residues essential for sensing BDSF. The effects of the BDSF treatment on the domain-deletion mutants with mutations in the coding sequences of the sensor or pairwise TMs included significant decreases in extracellular protease production (Fig. 3B and 3C), biofilm formation (Fig. 3D), and *engXCA* transcription (Fig. 3E). These findings suggested that the RpfC sensor and TM regions are crucial for detecting BDSF.

**FIG 3.**
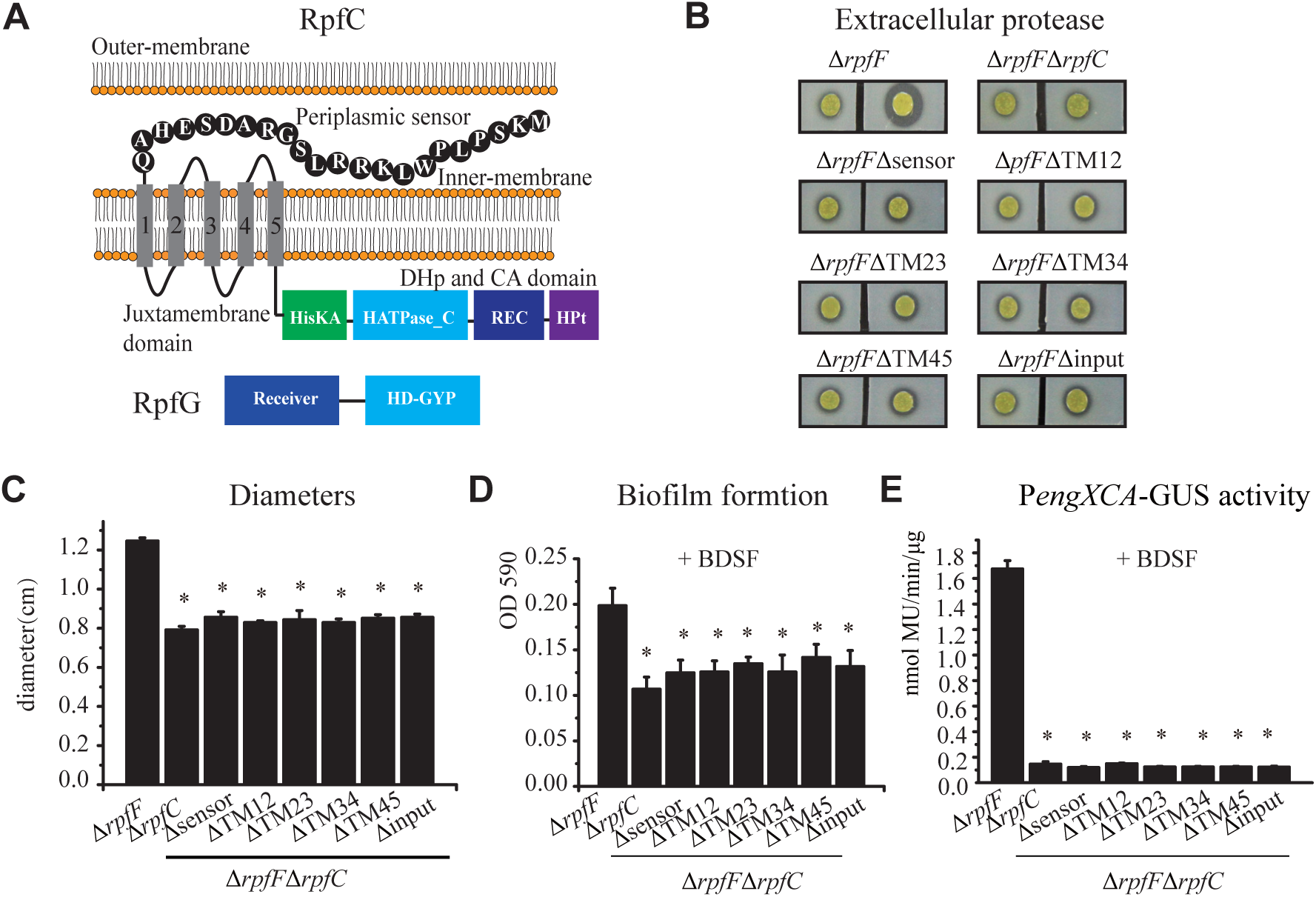
RpfC sensor and transmembrane regions are critical for sensing BDSF. (A) Schematic view of the full-length RpfC embedded in the membrane and RpfG. Secondary structures of RpfC-RpfG were predicted using Pfam. Transmembrane regions were predicted using TMpred. (B) Extracellular protease production assay. In each panel, *rpfF rpfC* double mutant (deletions in various regions) bacterial colonies on NYG medium are presented, with the effects of 4 μL BDSF added nearby the *rpfF rpfC* mutant to trigger quorum sensing presented on the right. (C) Protease activity was determined by measuring the diameters (cm) of the protein degradation zones after a 36-h incubation (in A and B, n = 4). (D) Biofilm production. A crystal violet staining method was used to quantify the biofilm formation of various *rpfF rpfC* mutants (n = 4). 200 μM BDSF was added to the bacterial culture. (E) P*engXCA* reporter activity assay to evaluate the effect of 50 μM BDSF. Each experiment was repeated four times. The vertical bar represents the standard deviation (n = 4). The asterisk indicates a significant difference compared with the unstimulated control (*rpfF* mutant).

Further analyses focused on identifying essential amino acid residues within the RpfC sensor. Regarding extracellular protease production, in the presence of 200 µM BDSF, seven point mutations in the RpfC sensor sequence (i.e., W7A, R15A, D17A, S18A, E19A, H20A, and Q22A) resulted in a significant decrease in extracellular protease production (Fig. 4A and 4B). For the biofilm quantification assay, the K2A, P4A, P6A, W7A, L8A, R10A, R11A, L12A, S13A, G14A, R15A, D17A, S18A, E19A, H20A, and Q22A substitutions were detrimental to biofilm formation (Fig. 4C). The P*engXCA*-*gus* reporter assay was again used to quantify transcriptional changes induced by the BDSF treatment. The L5A, W7A, R10A, L12A, R15A, D17A, S18A, E19A, H20A, and Q22A mutations resulted in significant decreases in *engXCA* transcription, whereas G14A had the opposite effect (Fig. 4D).

**FIG 4.**
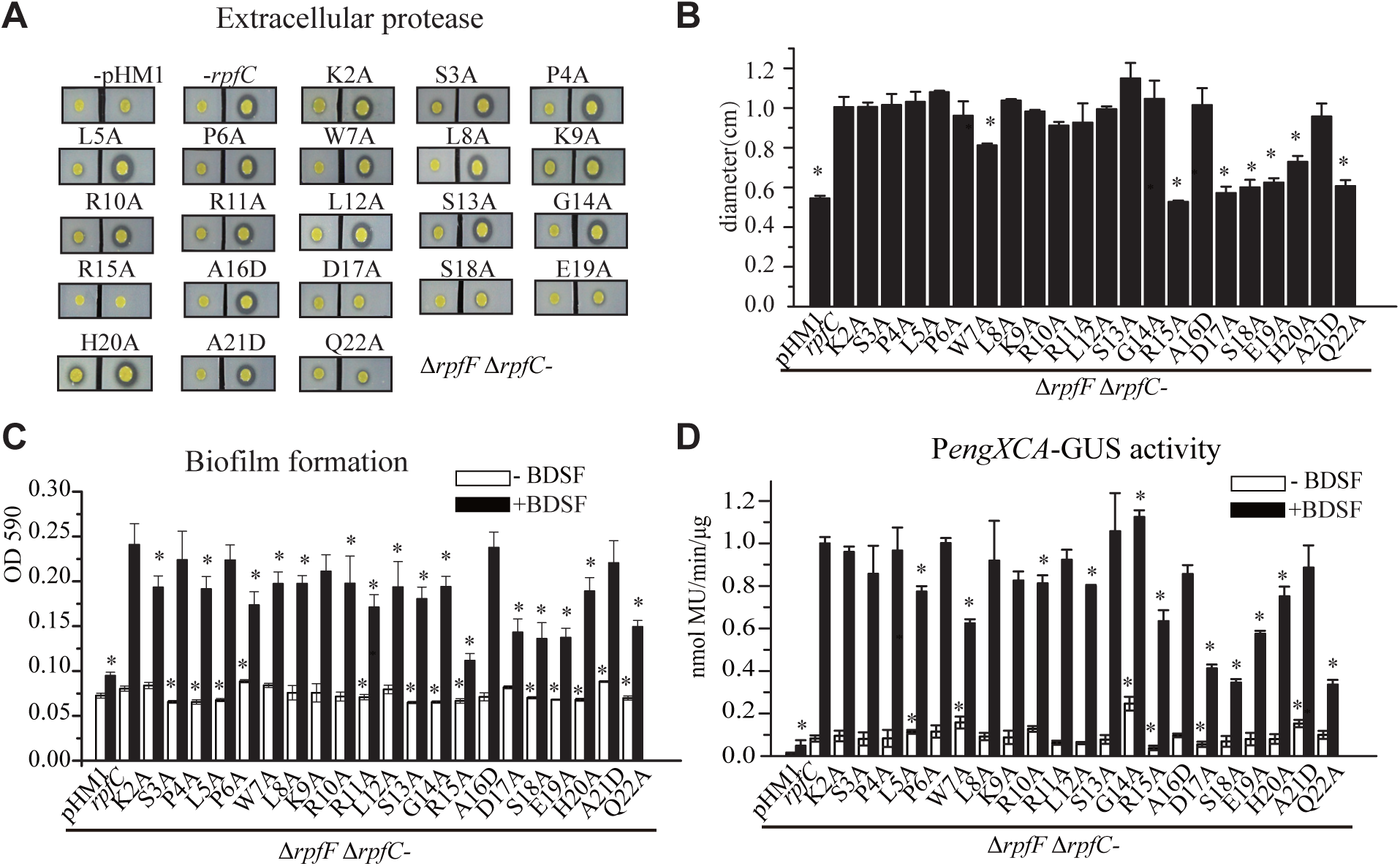
Identification of the amino acid residues in the RpfC sensor that are essential for sensing BDSF. *rpfC* mutants with point mutations generated by alanine-scanning mutagenesis in the *rpfF* background were tested. (A) Extracellular protease production by the mutants. In each panel, *rpfF rpfC* double mutant (various substitution sites) bacterial colonies on NYG medium are presented, with the effects of 4 μL BDSF added nearby the *rpfF rpfC* mutant to trigger quorum sensing presented on the right. (B) Protease activity was determined by measuring the diameters (cm) of the protein degradation zones after a 36-h incubation (in both A and B, n = 4). (C) Biofilm production by the mutants. A crystal violet staining method was used to quantify the biofilm formation of various mutants (n = 4). 50 μM BDSF was added to the bacterial culture. (D) P*engXCA* reporter activity assay to evaluate the effect of 50 μM BDSF. Each experiment was repeated four times. The vertical bar represents the standard deviation (n = 4). The asterisk indicates a significant difference compared with the unstimulated control (*rpfF* mutant).

Although point mutations in the RpfC sensor sequence have complex effects on diverse phenotypes, the above-mentioned results combined with the findings of an earlier study on the DSF–RpfC sensor interaction (7) revealed six amino acid residues in the RpfC ectodomain facing the inner-membrane region (i.e., R15A, D17A, S18A, E19A, H20A, and Q22A) are essential for ligand sensing or for maintaining the protein conformation. Substitutions of these residues significantly decreased BDSF- and DSF-triggered quorum-sensing signaling in *X. campestris* pv. *campestris*.

### High-affinity binding of BDSF to the RpfC sensor activates autokinase activity

Only full-length RpfC embedded in a liposome has enzymatic activity (Fig. S1). The autophosphorylation assay demonstrated that a low BDSF concentration (0.5 µM) induced the RpfC autokinase activity in a manner similar to that of DSF (Fig. 5A). Increases in the BDSF concentrations resulted in gradual increases in RpfC autophosphorylation levels (Fig. 5B). A dose–response analysis revealed that the half-maximal effective concentration (EC_50_) of BDSF for activating RpfC was approximately 0.06 µM (Fig. 5C), which is slightly lower than that of DSF (7). Regarding the ectodomain residues near the membrane, recombinant RpfC^R15A^, RpfC^D17A^, RpfC^E19A^, RpfC^H20A^, and RpfC^Q22A^ lacked autokinase activity (Fig. 5D), regardless of the presence of BDSF, implying that these residues are important for stabilizing the natural RpfC conformation. Notably, although RpfC^S18A^ lacked detectable autokinase activity in the absence of BDSF, the BDSF treatment resulted in increased autokinase activity (Fig. 5D), reflecting the importance of this residue for detecting BDSF. We also determined that a high *trans*-BDSF concentration can activate the RpfC autokinase, with an EC_50_ of 1.11 µM, which is 18 times the EC_50_ of BDSF (Fig. S2A and S2B).

**FIG 5.**
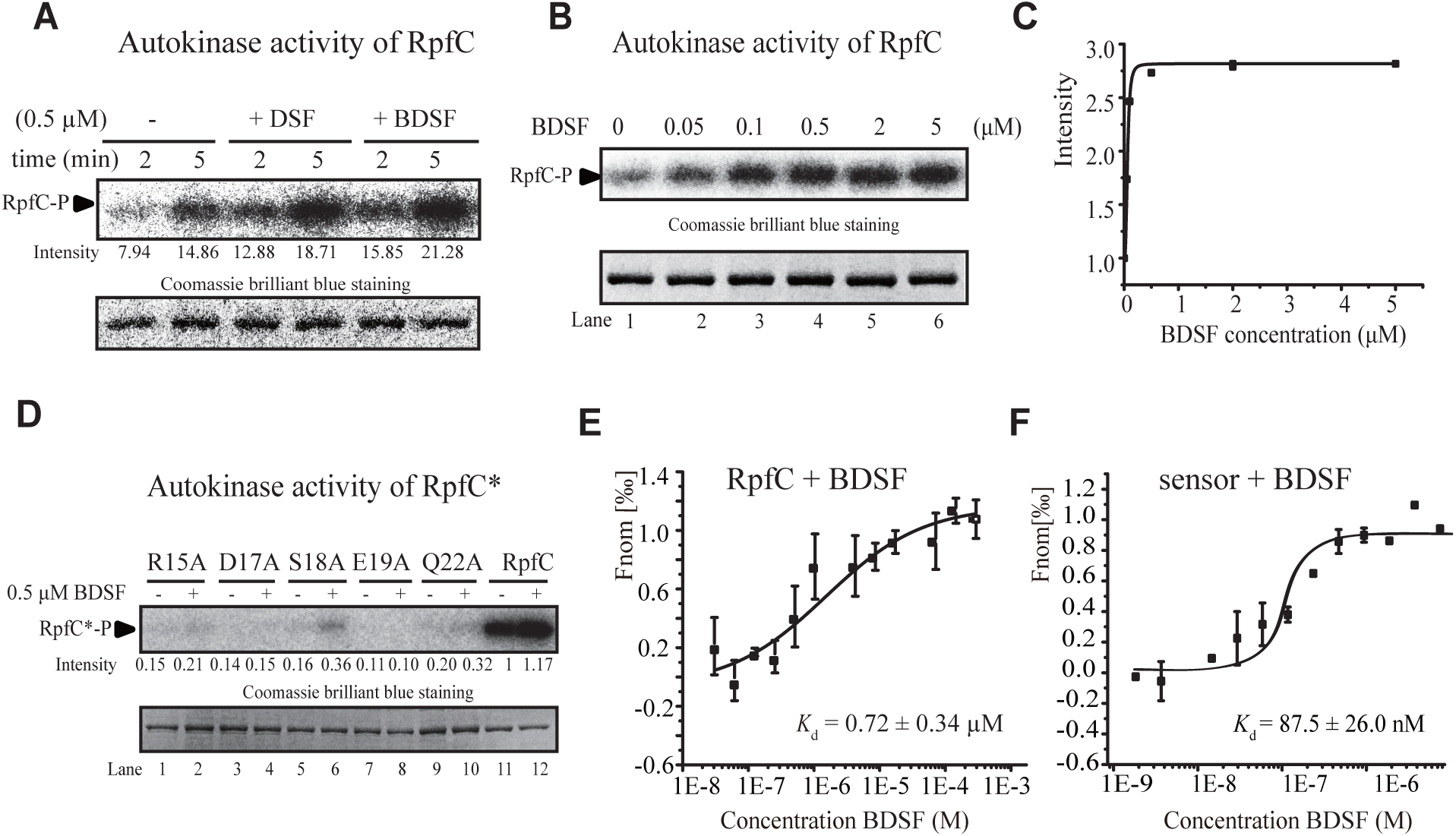
BDSF binds to the RpfC sensor to inactivate the autokinase. (A) BDSF and DSF similarly activate the RpfC autokinase. Lower panels present RpfC stained with Coomassie brilliant blue, which served as loading controls. (B and C) BDSF activates the RpfC autokinase in a dose-dependent manner. (B) Various concentrations of BDSF (0.05–5 μM) were added along with γ-[^32^P]ATP to the reaction mixture for the autophosphorylation assay. (C) Quantification of the isotopic signal for the RpfC autophosphorylation presented in (B). (D) Substitution of essential amino acids decreased the RpfC autophosphorylation level. (E and F) Microscale thermophoresis assay revealed that BDSF can bind to the full-length RpfC liposome (E) and RpfC sensor (F). The DSF concentrations ranged from 25 to 2,000 μM. The solid curve presents the fit of the data points to the standard KD-Fit function. Each binding assay was repeated three times and black bars represent the standard deviation. *K*_d_ = dissociation constant.

Previous research involving the microscale thermophoresis (MST) approach confirmed DSF can bind to the full-length RpfC and the RpfC sensor fragment, with a *K*_d_ of 140 ± 40 nM (7). Our MST assay indicated that BDSF can also bind to the full-length RpfC liposome, with a *K*_d_ of 0.72 ± 0.34 µM (Fig. 5E). However, BDSF did not bind to the truncated RpfC lacking the N-terminal sensor or input regions (Fig. S3A), suggesting that BDSF specifically binds to the RpfC sensor. Consequently, a peptide comprising the RpfC sensor was produced by fusing the corresponding coding sequence to a glutathione-S-transferase (GST) tag. The sensor was subsequently obtained via an on-column cleavage method that removed the GST tag. The MST assay was conducted to determine the affinity of the binding between BDSF and the RpfC sensor (*K*_d_ of 87.5 ± 26.0 nM) (Fig. 5F). As a negative control, BDSF did not bind to GST, suggesting that the observed BDSF–RpfC binding was not caused by residual GST (Fig. S3C). Additionally, recombinant RpfC^R15A^, RpfC^D17A^, RpfC^S18A^, RpfC^E19A^, RpfC^H20A^, and RpfC^Q22A^ were unable to bind to BDSF (Fig. S3D–3H), providing further evidence of the importance of these residues for maintaining the protein conformation required for sensing BDSF. These results indicated that BDSF binds to the RpfC sensor, with a binding affinity that is greater than that of the DSF–RpfC interaction.

The binding between *trans*-BDSF and the full-length RpfC liposome was also analyzed. The calculated *K*_d_ (11.22 ± 0.95 µM) (Fig. S2C) was substantially lower than the *K*_d_ for the BDSF–RpfC interaction.

### Neither BDSF nor DSF regulates RpfC phosphotransferase and phosphatase activities or the c-di-GMP degradation by the RpfG hydrolase

There were inconsistencies in the results of the physiological and biochemical analyses. For example, the physiological activity of BDSF for inducing quorum sensing was substantially lower than that of DSF (Fig. 1). However, the biochemical activity of BDSF was similar to that of DSF because BDSF bound to the RpfC sensor with a higher affinity than that of DSF and BDSF and DSF activated the RpfC autokinase to similar levels (Fig. 5). It was unclear which biochemical mechanism was responsible for the physiological difference between the two signals. In addition to autokinase activity, histidine kinases usually have phosphotransferase and phosphatase activities for phosphorylating and dephosphorylating cognate response regulators. The inconsistencies in the results of this study may indicate that BDSF regulates the phosphotransferase or phosphatase activity of RpfC toward the cognate RpfG.

There are no published reports describing the phosphorylation activity of the deduced RpfC–RpfG two-component signaling system. In a phosphotransferase assay using γ-[^32^P]ATP, the autophosphorylated full-length RpfC liposome transferred the phosphoryl group to RpfG (Fig. 6A). A substitution at the predicted phosphorylation site (Asp^80^) of RpfG prevented phosphorylation, demonstrating that RpfC phosphorylates RpfG at Asp^80^ (Fig. 6A). However, BDSF and DSF did not appear to affect the extent of the phosphorylation of RpfG (Fig.6B), implying neither BDSF nor DSF controls RpfC phosphotransferase activity.

**FIG 6.**
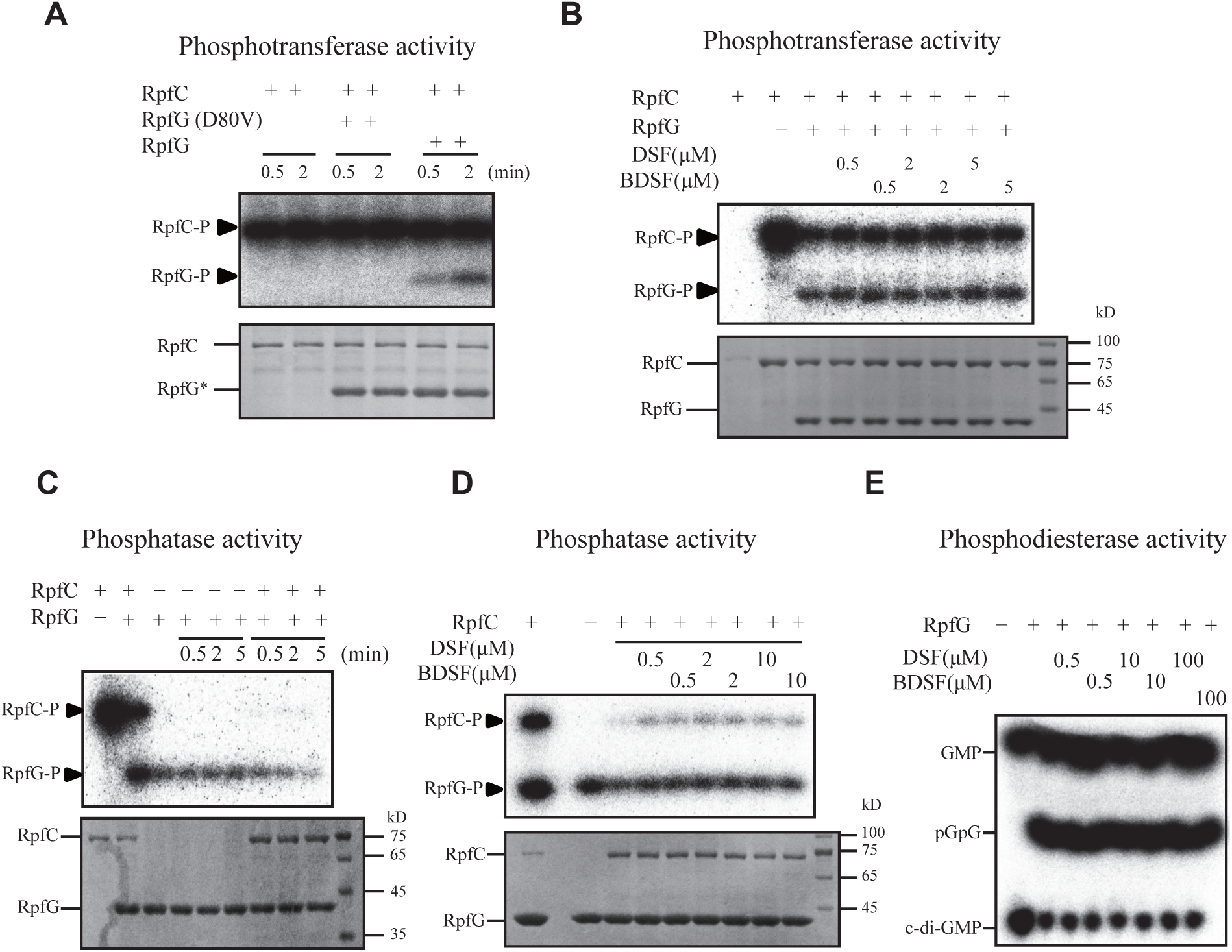
Both BDSF and DSF did not regulate RpfC phosphotransferase and phosphatase activities or the degradation of c-di-GMP by RpfG hydrolase. (A) RpfC phosphorylated RpfG at its Asp^80^ residue. The RpfC liposome was autophosphorylated by γ-[^32^P]ATP and recombinant RpfG or RpfG^D80V^ was added to the reaction mixtures. The phosphorylation signal was detected by autoradiography. Lower panels present proteins stained with Coomassie brilliant blue. (B) Addition of BDSF and DSF did not affect the RpfC-RpfG phosphotransfer. (C) Full-length RpfC can function as a phosphatase to dephosphorylate RpfG. After the RpfC-RpfG phosphotransfer, RpfC liposomes were removed by centrifugation and free ATP was removed by desalination. Fresh unphosphorylated RpfC liposomes were then added to the reaction mixture, which was then incubated for various periods. The phosphorylation signal was detected by autoradiography. (D) Addition of BDSF and DSF did not affect the RpfC phosphatase activity toward RpfG. (E) Addition of BDSF and DSF did not affect the degradation of c-di-GMP by RpfG hydrolase. The γ-[^32^P]-labeled c-di-GMP was synthesized using a recombinant tDGC cyclase and then co-incubated with recombinant RpfG. If necessary, BDSF and DSF were added to the reaction mixtures. The products of the enzymatic reaction were separated by thin layer chromatography and the isotopic signal was recorded by autoradiography. (A to E) Each experiment was repeated three times and representative results are presented.

To establish a phosphatase assay, after the RpfC–RpfG phosphotransfer, the phosphorylated RpfC liposome was discarded by high-speed centrifugation and γ-[^32^P]ATP was removed by desalination. The addition of fresh unphosphorylated RpfC liposomes to the reaction mixture decreased the RpfG phosphorylation level after a 2-min co-incubation (Fig. 6C), indicating that RpfC has a phosphatase that can dephosphorylate RpfG. However, the co-incubation with DSF or BDSF in the reaction mixture did not affect the RpfG phosphorylation level (Fig. 6D). Accordingly, BDSF and DSF likely do not control RpfC phosphatase activity.

The hydrolase activity of the RpfG C-terminal HD-GYP domain degrades c-di-GMP to GMP (Fig. 3A). Therefore, another possible explanation for the observed regulatory difference is that BDSF and DSF may differentially influence RpfG hydrolase activity. To test this hypothesis, we synthesized [^32^P]-labeled c-di-GMP using a recombinant cyclase with a modified GGDEF domain-containing cyclase tDGC (26). The degradation of c-di-GMP by the RpfG hydrolase was then examined after adding DSF or BDSF to the reaction mixture. The autoradiography results indicated that neither BDSF nor DSF affected the c-di-GMP degradation dynamics (Fig. 6E).

The results of these enzymatic analyses revealed that neither BDSF nor DSF regulates RpfC autokinase, phosphotransferase, and phosphatase activities or the RpfG hydrolase activity involved in degrading c-di-GMP. These factors did not contribute to the physiological differences between BDSF and DSF in triggering quorum sensing.

### A low BDSF concentration is enough to regulate quorum sensing in the *rpfB* mutant

In *Xanthomonas* spp., RpfB functions as a fatty acyl-CoA ligase that catalyzes the activation of fatty acids via a thioesterification with coenzyme A. The fatty acyl-CoA is then readily used for generating energy through β-oxidation (13, 14). Therefore, RpfB contributes to metabolism, especially the degradation of DSF-family signals. Because BDSF and DSF did not regulate RpfC–RpfG phosphorylation or RpfG hydrolase activity (Fig. 6), we speculated that the difference in RpfB-mediated degradation when BDSF or DSF was used as the substrate accounted for the physiological variation. To test this hypothesis, we constructed the *rpfF rpfB* double mutant (Δ*rpfF*–Δ*rpfB*), which cannot synthesize and degrade DSF-family signals, to functionally characterize RpfB during BDSF- and DSF-triggered quorum sensing. Regarding the *rpfF* mutant, which cannot biosynthesize DSF-family signals, the addition of 10 µM DSF significantly enhanced biofilm formation (i.e., approximately 2.6 times that induced by 10 µM BDSF) (Fig. 7A). In the Δ*rpfF*–Δ*rpfB* double mutant, in which exogenously applied BDSF and DSF cannot be degraded, the 10 µM DSF treatment significantly increased biofilm formation (i.e., 1.2 times that of the *rpfF* mutant). Moreover, the 10 µM BDSF treatment significantly enhanced the biofilm formation of the Δ*rpfF*–Δ*rpfB* double mutant to a level similar to that induced by 10 µM DSF and 3.0 times that of the *rpfF* mutant (Fig. 7A). The overexpression of *rpfB* in the double mutant restored the biofilm production level to that of the *rpfF* mutant (Fig. 7A). Furthermore, the *engXCA* transcription level induced by 10 µM BDSF was significantly lower than that induced by 10 µM DSF (approximately 0.23 times lower, Fig.7B). However, in the Δ*rpfF*–Δ*rpfB* double mutant, the *engXCA* transcription level induced by 10 µM BDSF was higher than that induced by 10 µM DSF (Fig. 7B). The overexpression of full-length *rpfB* in the double mutant significantly decreased *engXCA* transcription to a level similar to that in the *rpfF* mutant, suggesting *rpfB* encodes an indispensable regulator of this process (Fig. 7B).

**FIG 7.**
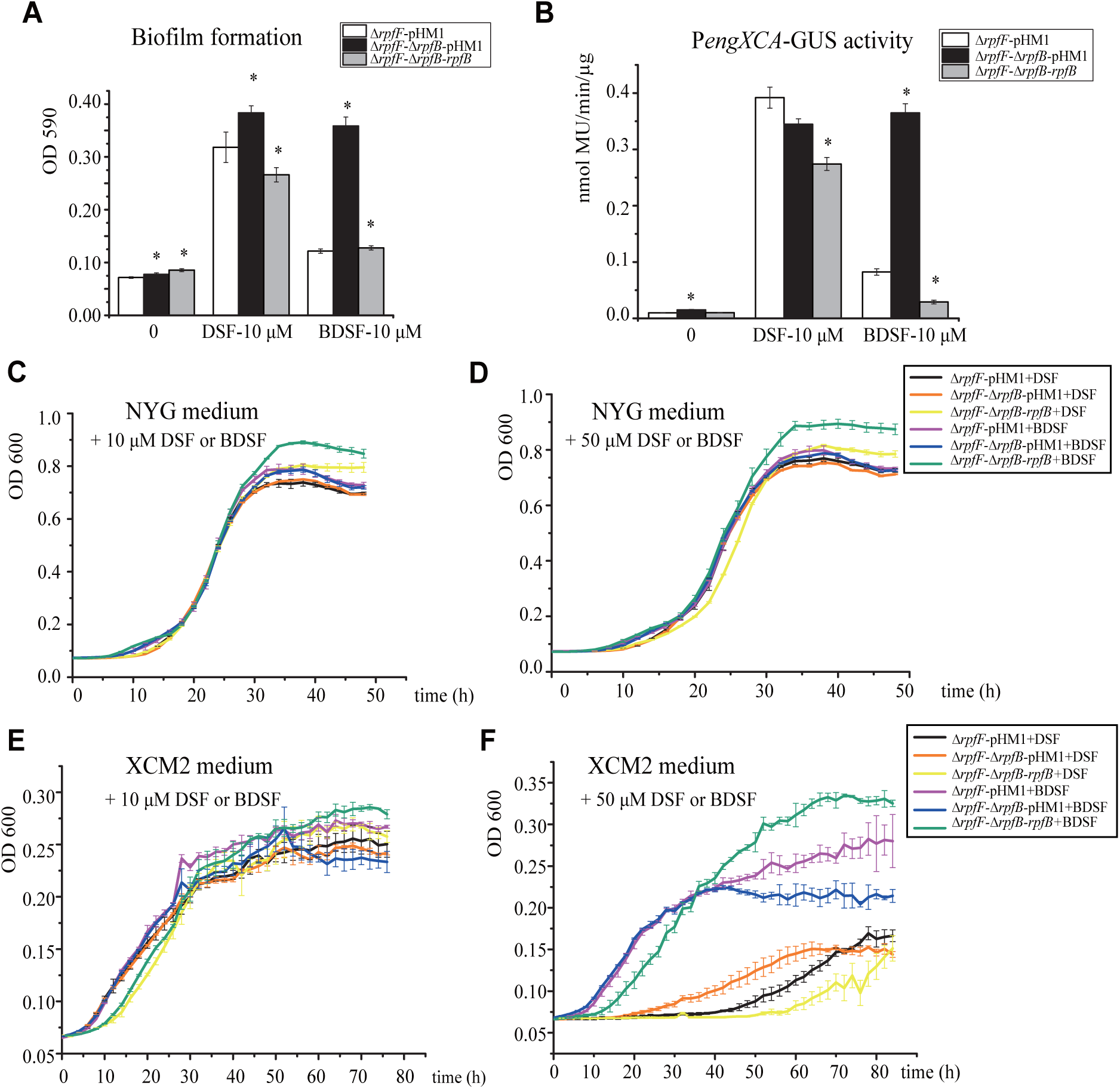
BDSF is prone to degradation in bacterial cells. (A) Quantification of the biofilm production in the *rpfF* mutant and the *rpfF rpfB* double mutant. 10 μM BDSF and DSF were added to the bacterial cultures. (B) P*engXCA* reporter activity assay of the *rpfF* mutant and the *rpfF rpfB* double mutant. 10 μM BDSF and DSF were added to the bacterial cultures to induce *engXCA* transcription. (C to F) Bacterial growth curves. Various strains were grown in rich medium (i.e., NYG) (C and D) and the type III secretion system-inducing medium (i.e., XCM2) (E and F). If necessary, DSF or BDSF was added. (A to F) Each experiment was repeated four times. The vertical bar represents the standard deviation (n = 4). The asterisk indicates a significant difference compared with the unstimulated control.

To investigate the consequences of the inability to degrade BDSF on bacteria, we analyzed bacterial growth under various conditions. In rich medium (i.e., NYG medium), there was no substantial difference in the growth of the *rpfF* mutant and the Δ*rpfF*–Δ*rpfB* double mutant, regardless of the presence of DSF or BDSF (Fig. 7C and 7D). In the XCM2 medium, which is usually used to mimic plant conditions to induce the type III secretion system, the Δ*rpfF*–Δ*rpfB* double mutant grew slightly faster than the *rpfF* mutant in the presence of 50 µM DSF (Fig. 7E), indicating that DSF at this concentration did not have a substantial effect on bacterial growth. However, in XCM2 medium supplemented with 50 µM BDSF, the Δ*rpfF*–Δ*rpfB* double mutant grew much slower than the *rpfF* mutant; the genetic complementation of *rpfB* in the double mutant significantly restored bacterial growth (Fig. 7F). Hence, the degradation of BDSF by RpfB was important for bacterial growth in the XCM2 medium.

The results revealed that RpfB substantially suppresses the physiological activity of BDSF during the induction of quorum sensing. However, BDSF and DSF activities are similar in the *rpfB* mutant, suggesting that the RpfB-mediated degradation of BDSF is the major determinant of the differential functions of BDSF and DSF in bacterial cells.

## DISCUSSIONS

The DSF-family chemicals are important quorum-sensing signals in Gram-negative bacteria and participate in the interkingdom communication between bacteria and fungi (5, 27, 28). Various DSF-family chemicals have been identified in *Xanthomonas*, *Pseudomonas*, *Stenotrophomonas*, and *Burkholderia* bacterial species, including DSF, BDSF, IDSF, CDSF, and SDSF (10, 29). These chemicals are structurally similar, but their abundance, precursors, cellular dynamics, and functions have not been thoroughly investigated. The only chemical difference between BDSF and DSF is the lack of a methyl group at the C-11 site of BDSF (Fig. 1A). In this study, we demonstrated that compared with DSF, BDSF is a less active quorum-sensing signal. Moreover, *trans*-BDSF does not appear to have any major physiological effects in *X. campestris* pv. *campestris*. High BDSF concentrations were needed to trigger quorum sensing-modulated processes, including the production of cellulase and extracellular proteases as well as biofilm development (Fig. 1). However, an interesting inconsistency was detected in the biochemical activity of BDSF. Similar to DSF (7), BDSF is detected by a sensor histidine kinase in *X. campestris* pv. *campestris* (Fig. 3 and Fig. 4), but BDSF binds to the RpfC sensor region with an affinity higher than that of DSF, and 0.5 µM DSF and BDSF activated RpfC autophosphorylation to similar levels (Fig. 5). This apparent discrepancy between the physiological and biochemical activities of BDSF compelled us to clarify the molecular basis of the low physiological activity of BDSF. We revealed that BDSF is easily degraded by RpfB, which is the main reason for the low physiological activity of BDSF, rather than the RpfC phosphotransfer or phosphatase activity toward the cognate response regulator RpfG or the RpfG hydrolase-mediated degradation of the bacterial second messenger c-di-GMP (Fig. 6 and Fig. 7). In the *rpfB* mutant, a low BDSF concentration was sufficient for regulating quorum sensing at the same level as DSF (Fig. 7). During the subsequent analysis, a high BDSF concentration arrested bacterial growth under conditions that mimic those in plants. Therefore, *X. campestris* pv. *campestris* tightly regulates the BDSF concentration and activity, especially in host plants.

In *Xanthomonas* species, the sensor histidine kinase RpfC was predicted to be the receptor of DSF-family signals (1), but this was not experimentally verified until the DSF– RpfC interaction was confirmed (7). The issues that contributed to the delayed confirmation of this interaction included the fact that unlike RpfR, RpfC is a membrane-bound protein with five putative TM helices (Fig. 2A). A soluble truncated RpfC protein without sensor or TM regions does not have any autokinase activity (Fig. S1). Therefore, full-length RpfC must be embedded in membranes [e.g., inverted membrane vesicle (IMV), liposome, and nanodiscs] to fold into the conformation required for enzymatic activities. Additionally, protein–fatty acid interactions are difficult to detect using available methods (30, 31) involving surface plasmon resonance, biolayer interferometry, and isothermal titration calorimetry. In the present study, we constructed the RpfC liposome and established biochemical assays for analyzing autokinase, phosphotransferase, and phosphatase activities to characterize RpfC– RpfG phosphorylation (Fig. 6). The MST assay revealed that BDSF binds to the RpfC sensor and then activates RpfC autophosphorylation similar to DSF. The essential residues of the RpfC sensor region, especially those near the inner-membrane region, are critical for maintaining the RpfC structure or sensing DSF and BDSF. Therefore, RpfC is the receptor of BDSF and DSF. The similarity in their structures suggests that RpfC is also the receptor of other DSF-family signals, but the precise effects of these signals on RpfC remain to be investigated.

Structurally, BDSF and DSF are similar to farnesol, which controls the life stage transitions and biofilm development of *C. albicans* (32). In *C. albicans*, low concentrations of the two BDSF isomers (e.g., 3 µM) can significantly inhibit biofilm formation more effectively than farnesol. At high concentrations (e.g., 30–300 µM), BDSF and *trans*-BDSF can arrest the yeast-to-hypha conversion and growth (15, 16). Therefore, both BDSF and *trans*-BDSF have biological effects on *C. albicans*, making them potentially useful for treating infections caused by this fungus. However, we observed that the addition of a high *trans*-BDSF concentration failed to stimulate the extracellular protease production and biofilm formation of *X. campestris* pv. *campestris* (Fig. 2A and 2B). At 100 µM, the *engXCA* transcription level induced by BDSF was 4.1 times that induced by *trans*-BDSF. Moreover, *trans*-BDSF concentrations are never as high as 100 µM in bacterial cells. Thus, *trans*-BDSF seems not a natural quorum-sensing signal of *X. campestris* pv. *campestris*. The MST assay data indicated that *trans*-BDSF can bind to the RpfC liposome, but at an affinity (*K*_d_ of 11.22 ± 0.95 µM) that is lower than that for BDSF and DSF. Consistent with this binding affinity, the EC_50_ of *trans*-BDSF was 18 times that of BDSF and DSF according to the autokinase assay. These findings provide further evidence of the low-affinity binding and relatively low induction of RpfC autophosphorylation that prevents *trans*-BDSF from being a quorum-sensing signal for *X. campestris* pv. *campestris*. Therefore, the *X. campestris* pv. *campestris* RpfC can discriminate between BDSF and *trans*-BDSF, but the corresponding *C. albicans* receptor, which remains unknown, cannot.

This study revealed that though BDSF is a less active signaling molecule than DSF, both compounds have similar biochemical activities. More specifically, they bind to the RpfC sensor region with a similar affinity and their EC_50_ for inducing RpfC autokinase activity are also similar. We eliminated the possibility that the observed physiological difference was caused by BDSF/DSF-triggered changes in RpfC phosphotransferase and phosphatase activities toward RpfG or the ability of RpfG hydrolase to degrade c-di-GMP. Both BDSF and DSF did not affect these enzymatic processes. However, in the *rpfB* mutant, which cannot degrade DSF-family molecules, a low BDSF concentration can induce quorum-sensing responses at the same level as DSF. In wild-type strains, the BDSF concentration must be approximately 5 times that of DSF to induce similar quorum-sensing activities. Therefore, BDSF is more likely to be metabolized in bacterial cells than DSF. Considering that BDSF is the most abundant DSF-family compound when *X. campestris* pv. *campestris* is grown in host plants or on medium containing plant extracts, the ease with which BDSF can be degraded may have important adaptive consequences for bacterial infection or growth. Our phenotypic characterization revealed that in rich medium, high concentrations of DSF or BDSF did not substantially modulate bacterial growth. However, in the XCM2 medium used to simulate plant conditions and induce the bacterial type III secretion system, high BDSF concentrations have an inhibitory effect on bacterial growth (33, 34), whereas DSF at the same concentrations does not affect growth. During an infection of host plants by *X. campestris* pv. *campestris*, BDSF serves as the major quorum-sensing signal, but because its accumulation adversely affects bacterial growth, RpfB is required to decrease its concentration. Therefore, *X. campestris* pv. *campestris* evolved a molecular mechanism to tightly regulate the biosynthesis and degradation of BDSF in host plants, which differs from the regulatory control associated with its analog DSF.

## ACKONOWLEGEMENT

We thank members of our laboratory for helping discussion. This work was supported by National Natural Science Foundation of China (31825021), CAS Center for Excellence in Biotic Interactions and State Key Laboratory of Plant Genomics.

We declare no conflict of interest relevant to the studies reported.

## MATERIALS AND METHODS

### Bacterial strains and plasmids

All bacterial strains and recombinant vectors used in this study are listed in Table S1. *Xanthomonas campestris* pv. *campestris* 8004-derived strains and the wild-type strain were grown at 28 °C in NYG medium (5 g L^−1^ tryptone, 3 g L^−1^ yeast extract, and 20 g L^−1^ glycerol, pH 7.0) or 210 medium (5 g L^−1^ sucrose, 8 g L^−1^ casein enzymatic hydrolysate, 4 g L^−1^ yeast extract, 3 g L^−1^ K_2_HPO_4_, and 0.3 g L^−1^ MgSO_4_·7H_2_O, pH 7.0). Growth curve measuments were conducted in XCM2 medium (2.36 g L^−1^ succinic acid, 0.15 g L^−1^ casamino acids, 1g L^−1^ [NH_4_]_2_SO_4_, 0.01 mM MgSO_4_, 13.8 g L^−1^ K_2_HPO_4_.3H_2_O, 8.35 g L^−1^ KH_2_PO_4_). *Escherichia coli* DH5α was used as the host for constructing all recombinant vectors, whereas *E. coli* BL21 (DE3) was used as the host for expressing recombinant proteins encoded in the pET30a vector (Novagen, USA). Appropriate antibiotics were added when needed at the following concentrations: kanamycin (50 µg mL^−1^), spectinomycin (150 µg mL^−1^), ampicillin (100 µg mL^−1^), and rifamycin (25 µg mL^−1^). *Xanthomonas campestris* pv. *campestris* 8004 and *E. coli* electrocompetent cells were prepared by extensively washing bacterial cells three times with ice-cold 10% glycerol. Both *X. campestris* pv. *campestris* and *E. coli* cells were transformed using the Pulser XCell system (Bio-Rad, USA), with the following conditions: 1.8 kV cm^−1^, 25 µF, and 200 Ω. The following HPLC-purified DSFs were used in this study: DSF (CAS No. 677354-23-3, purity >90.0%) and BDSF (CAS No. 55928-65-9, purity >95.0%) purchased from Sigma Aldrich (USA) and *trans*-BDSF (CAS No. 4412-16-2, purity >95.0%) purchased from WuXi AppTec (China). The chemicals were diluted to the concentrations required for the various experiments.

### Mutant construction and genetic complementation

Unless specified otherwise, standard molecular biology techniques (e.g., PCR, DNA ligation, restriction enzyme digestion, and western blotting) were performed according to the protocols in Molecular Cloning. The in-frame deletion (markerless) mutants *rpfB* and *rpfF rpfB* (Δ*rpfF*–Δ*rpfB*) were constructed using the suicide vector pK18mobsacB according to a homologous, double crossover method. Briefly, the 5′ and 3′ genomic sequences of a targeted region were amplified using primers listed in Table S2. The correct PCR products were ligated to pK18mobsacB (35). The recombinant pK18mobsacB vector was inserted into *X. campestris* pv. *campestris* 8004 competent cells via electroporation to generate single-crossover mutants that were selected in plates containing NYG medium supplemented with kanamycin. The single-crossover mutants were subsequently cultured in NYG medium (antibiotic free) for 1– 2 h and then grown in plates containing NYG medium supplemented with 10% sucrose to screen for second-round homologous cross-overs. Candidate bacterial mutants (i.e., resistant to 10% sucrose but sensitive to kanamycin) were verified by PCR and sequencing. To genetically complement the Δ*rpfB* mutant, the full-length *rpfB* sequence was amplified by PCR using primers listed in Table S2 and then ligated to the broad-host vector pHM1 . The recombinant vector was inserted into *E. coli* DH5α cells via electroporation. The vector was then extracted from *E. coli* DH5α cells and inserted into the Δ*rpfB* or Δ*rpfB–*Δ*rpfF* mutant, in which the transcription of the full-length *rpfB* sequence was under the control of a PlacZ promoter. Additionally, the Δ*rpfF*, Δ*rpfC*, and Δ*rpfC* Δ*rpfF* mutants and these strains with point mutations as well as the P*engXCA*-*gus* reporter strains and the RpfC and RpfC point mutation-protein expression strains were generated in a previous study .

### GUS activity assay

The GUS activity assay was performed as previously described (7). Bacterial strains were cultured and adjusted to an optical density at 600 nm (OD_600_) of 0.1, after which they were treated with DSF/BDSF/*trans*-BDSF for about 8 h. Cells were collected by centrifugation (12,000 *g* for 10 min at 4 °C) and immediately frozen in liquid nitrogen. The cells were resuspended in GUS extraction buffer (50 mM sodium phosphate, pH 7.0, 5 mM DTT, and 1 mM EDTA, pH 8.0) and then lysed by sonication. The mixture was centrifuged (12,000 *g* for 10 min at 4 °C) and the supernatant was collected for the GUS activity assay, which was performed using 4-methylumbelliferyl-β-D-glucuronide (4-MUG; Sigma Aldrich) as a substrate. A standard curve was prepared by diluting the 4-MUG stock solution. The fluorescence of the samples and solutions used to prepare standard curves was measured using excitation and emission wavelengths of 360 and 460 nm, respectively. The supernatant protein concentrations were determined using the Coomassie brilliant blue G-250 Protein Assay kit (Bio-Rad), with BSA serving as the standard. Each experiment was performed with at least three independent replicates.

### Biofilm formation assay

The biofilm formation assay was conducted using crystal violet as previously described (36). Bacterial strains were cultured and adjusted to an OD_600_ of 0.1 before adding an appropriate concentration of DSF/BDSF/*trans*-BDSF and then culturing for approximately 8 h at 28 °C. A 200-µL aliquot of the culture was added to a 96-well plate (Costar, USA) and incubated overnight. Bacterial growth in the 96-well plate was analyzed on the basis of OD_600_ values determined using the Infinite 200 Pro microplate reader (Tecan). The wells were washed with water before adding 1% crystal violet and incubating for 20 min. After the wells were washed with water, the crystal violet stain was solubilized in absolute ethanol and then biofilm formation was quantified according to the OD_590_ values determined using a microplate reader.

### Extracellular protease assay

The extracellular protease assay was conducted using NYG-milk medium in plates as previously described (37). If necessary, 3.5 µL DSF/BDSF/*trans*-BDSF was added to the plates (near the bacterial colony).

### Expression and purification of recombinant proteins

Poly-histidine (His_6_)-tagged proteins were expressed and purified by affinity chromatography using Ni-NTA agarose beads (Novagen, USA) according to the manufacturer-recommended instructions. Purified proteins were concentrated using Centricon YM-10 columns (Millipore) and the eluants were replaced by storage buffer (50 mM Tris-HCl, pH 8.0, 0.5 mM EDTA, 50 mM NaCl, and 5% glycerol).

### Reconstruction of membrane-bound histidine kinases using liposomes

Liposomes were reconstituted as previously described (7). Briefly, RpfC-containing IMVs were obtained as described above and then dissolved in suspension buffer (20 mM phosphate, 500 mM NaCl, and 20 mM imidazole, pH 7.4) for a final concentration of 10 mg mL^−1^. To prepare liposomes, 900 µL IMV suspension was mixed (end-over-end) with 100 µL 10% n-dodecyl-β-D-maltoside (DDM) for 45 min at 4 °C. The supernatant was collected by centrifugation (50,000 *g* for 30 min) and then used for a Ni-affinity chromatographic purification. The Ni-NTA beads (Novagen, USA) were pre-equilibrated with 5 volumes of binding buffer (20 mM phosphate, 500 mM NaCl, 20 mM imidazole, pH 7.4, and 0.1% DDM). The solubilized IMV and pre-equilibrated beads were mixed and incubated at 4 °C for 30 min. The supernatant was removed and the pellet was washed with binding buffer until the absorbance at OD_280_ returned to baseline. Finally, 100 µL elution buffer (20 mM phosphate, 500 mM NaCl, 250 mM imidazole, pH 7.4, and 0.1% DDM) was added to elute the purified RpfC–His6. To embed RpfC in liposomes, 10 mg liposomes (Avanti Polar Lipids) were dissolved in 1 mL buffer (50 mM Tris-HCl, pH 7.5, 10% glycerol, and 0.47% Triton-100). The purified RpfC–His6 in elution buffer was then added and the mixture was stirred at 4 °C for 45 min. The final ratio of phospholipids to proteins was about 10:1 (w/w). Bio-Beads (beads:detergent = 10:1; Bio-Rad) were added to remove the detergent and the solution was stirred gently at 4 °C overnight. Residual detergent was removed completely by adding Bio-Beads and then incubating for an additional 2 h. The Bio-Beads were removed using pipettes and then the liposomes containing RpfC–His6 were collected by centrifugation (200,000 *g* for 30 min at 4 °C). The RpfC liposomes were resuspended in the final buffer (20 mM Tris-HCl, pH 7.5, and 10% glycerol) and then stored at −80 °C until used.

### *In vitro* phosphorylation assay

*In vitro* phosphorylation and dephosphorylation assays were conducted as previously described (38). To assess autophosphorylation, an appropriate concentration of DSF/BDSF/*trans*-BDSF was mixed with the RpfC liposomes and the resulting solution was incubated with 100 µM ATP containing 10 µCi [γ-^32^P]ATP (PerkinElmer, USA) in autophosphorylation buffer (50 mM Tris-HCl, pH 7.8, 2 mM DTT, 25 mM NaCl, 25 mM KCl, and 5 mM MgCl_2_) at 28 °C for specific periods. For the phosphotransferase assay, RpfC was first autophosphorylated for 10 min to accumulate enough phosphorylated RpfC. If necessary, RpfC was removed after a high-speed centrifugation. Next, RpfG or RpfG mixed with DSF/BDSF/*trans*-BDSF was added to the reaction mixture. The phosphatase assay was conducted on the basis of the phosphotransferase assay. Specifically, [γ-^32^P]ATP was removed using a desalination column (PD-Sprintrap G-25). If necessary, DSF/BDSF/*trans*-BDSF was incubated with phosphorylated RpfG for 10 min. Fresh RpfC liposomes were added to the above reaction mixture to examine dephosphorylation. Each reaction was stopped by adding 5× SDS-PAGE loading buffer. The phosphorylated proteins were separated by 12% SDS-PAGE. The gels were placed in a Ziploc bag and exposed to a phosphor screen for 1 h. The screen was scanned using the PhosphorImage system (Amersham Biosciences, USA). If necessary, the signal intensity was quantified using ImageJ software.

### Purification and on-column cleavage of the sensor–GST fusion protein

The GST-tagged RpfC sensor was obtained and purified using GST Resin (TRANS) according to the GST Gene Fusion System Handbook (Amersham Biosciences). The GST tag was removed from the purified RpfC sensor during an on-column cleavage procedure. Briefly, the lysate of the recombinant *E. coli* BL21 (DE3) strain was mixed with pre-equilibrated GST Resin in PBS buffer for 10 min before being loaded on the column, which was washed using PBS buffer and equilibrated using PreScission buffer (50 mM Tris-HCl, pH 8.5, and 150 mM NaCl). Following the injection of 2 units PreScission Protease (GenScript), the column was sealed and placed on a rotator at 4 °C. After a 10-h digestion, the flowthrough fractions, which contained the sensor peptide without the GST tag, were collected. If necessary, the sensor peptide was purified by size exclusion chromatography using the Superdex 75 10/300 GL column (GE Healthcare) and then stored at −80 °C before use.

### MST assay

The binding of RpfC to BDSF or *trans*-BDSF was analyzed in an MST assay conducted using the Monolith NT.LabelFree instrument (Nano Temper Technologies GMBH, Germany), which can detect changes in size, charge, and conformation induced by binding. For this assay, RpfC liposomes were collected by centrifugation (200,000 *g* for 40 min) and then resuspended in MST buffer (50 mM Tris-HCl, pH 7.8, 150 mM NaCl, 10 mM MgCl_2_, and 0.1% Pluronic F-127) for a final concentration of approximately 0.1 µM. A range of BDSF concentrations (0.06–500 µM) in assay buffer (50 mM Tris-HCl, pH 7.8, 150 mM NaCl, 10 mM MgCl_2_, 0.05% Tween-20, and 0.5% methanol) was added to the RpfC liposome solution (1:1, v/v). After a 10-min incubation, the sample was loaded into NT.LabelFree standard capillaries and analyzed using 20% LED power and high MST power.

The purified sensor protein was dissolved in reaction buffer (50 mM Tris-HCl, pH 8.5, 150 mM NaCl, and 0.1% Pluronic F-127) for a final concentration of 0.1 µM. Diluted BDSF (4.58 × 10^−4^ µM to 15 µM) in buffer (50 mM Tris-HCl, pH 8.5, 150 mM NaCl, and 0.15‰ methanol) was used to prepare different BDSF concentrations, which were mixed with the sensor protein (1:1, v/v) and then loaded into NT.LabelFree standard capillaries. The label-free MST assay was performed using 20% LED power and high MST power. The KD Fit function of the Nano Temper Analysis Software (version 1.5.41) was used to calculate *K*_d_.

### Synthesis of [^32^P]c-di-GMP for the phosphodiesterase activity assay

The [^32^P]-labeled c-di-GMP was chemically synthesized using labeled [^32^P]GTP (3000 Ci/mmol; PerkinElmer, USA) and the purified His6-tagged enzyme tDGC (26), which is the *Thermotoga maritima* diguanylate cyclase with a key residue mutation (R158A) . A mixture comprising 5 µM tDGC and 20 µCi [^32^P]GTP (with 50 µM cold GTP) was added to 20 µL reaction buffer (300 mM NaCl, 50 mM Tris-HCl, pH 7.5, 20 mM MgCl_2_, and 2 mM DTT). After a 3-h incubation at 45 °C, the reaction was terminated by heating at 98 °C for 10 min. The precipitated protein was removed by centrifugation (20,000 *g* for 5 min). Radioactive [^32^P]c-di-GMP was detected on a polyethyleneimine-cellulose plate [1:1.5 (v/v) saturated (NH_4_)_2_SO_4_ and 1.5 M KH_2_PO_4_, pH 3.6]. Without any further purification, the mixture contained more than 95% [^32^P]c-di GMP.

The phosphodiesterase activity assay was performed as previously described (39). Purified RpfG was added to the reaction buffer consisting of 50 mM Tris-HCl, pH 7.5, 250 mM NaCl, 25 mM KCl, 10 mM MgCl_2_, and 2 mM DTT. Reactions were initiated by adding approximately 1 µM [^32^P]c-di-GMP substrate. Samples were incubated at 28 °C for 10 min before terminating the reactions by adding an equal volume of 0.5 M EDTA, pH 8.0. The reaction products were mixed with an equal volume of running buffer comprising 1:1.5 (v/v) saturated (NH_4_)_2_SO_4_ and 1.5 M KH_2_PO_4_, pH 3.6. A 2-µL aliquot of the reaction mixture was spotted and dried on Cellulose PEI TLC plates (Selecto Scientific, USA), which were subsequently developed in running buffer, air-dried, and exposed to a storage phosphor screen (GE Healthcare). The autoradiographic signals were detected and recorded using the Typhoon FLA7000 system (GE Healthcare).

### Bacteria growth curve assays

Bactaria growth curves were measured in NYG medium and XCM2 medium. Strains including Δ*rpfF*-pHM1, Δ*rpfF*-Δ*rpfB*-pHM1 and Δ*rpfF*-Δ*rpfB-rpfB* were cultured overnight and washed three times with XCM2 medium. Then they were adjusted to an optical density at 600 nm (OD_600_) of 0.4. Under NYG medium, bacteria solution was diluted 1000 times; under XCM2 medium, bacteria solution was diluted 100 times. In each condition, 10 µM DSF or BDSF and 50 µM DSF or BDSF was added into the culture. Growth curves were obtained after 2-4 days’ cultivation.

## Supplementary Information

**Supplementary Table 1.**
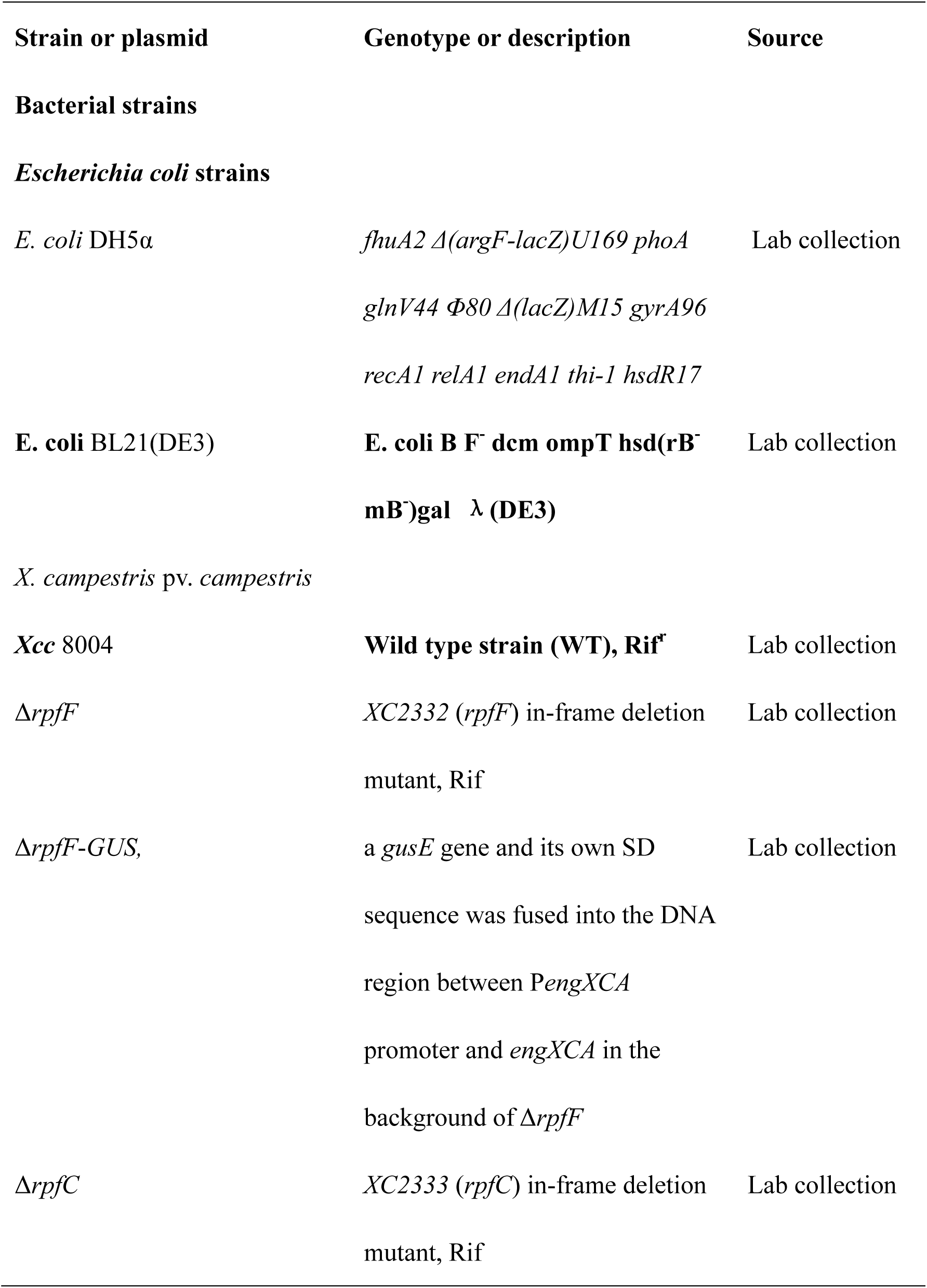
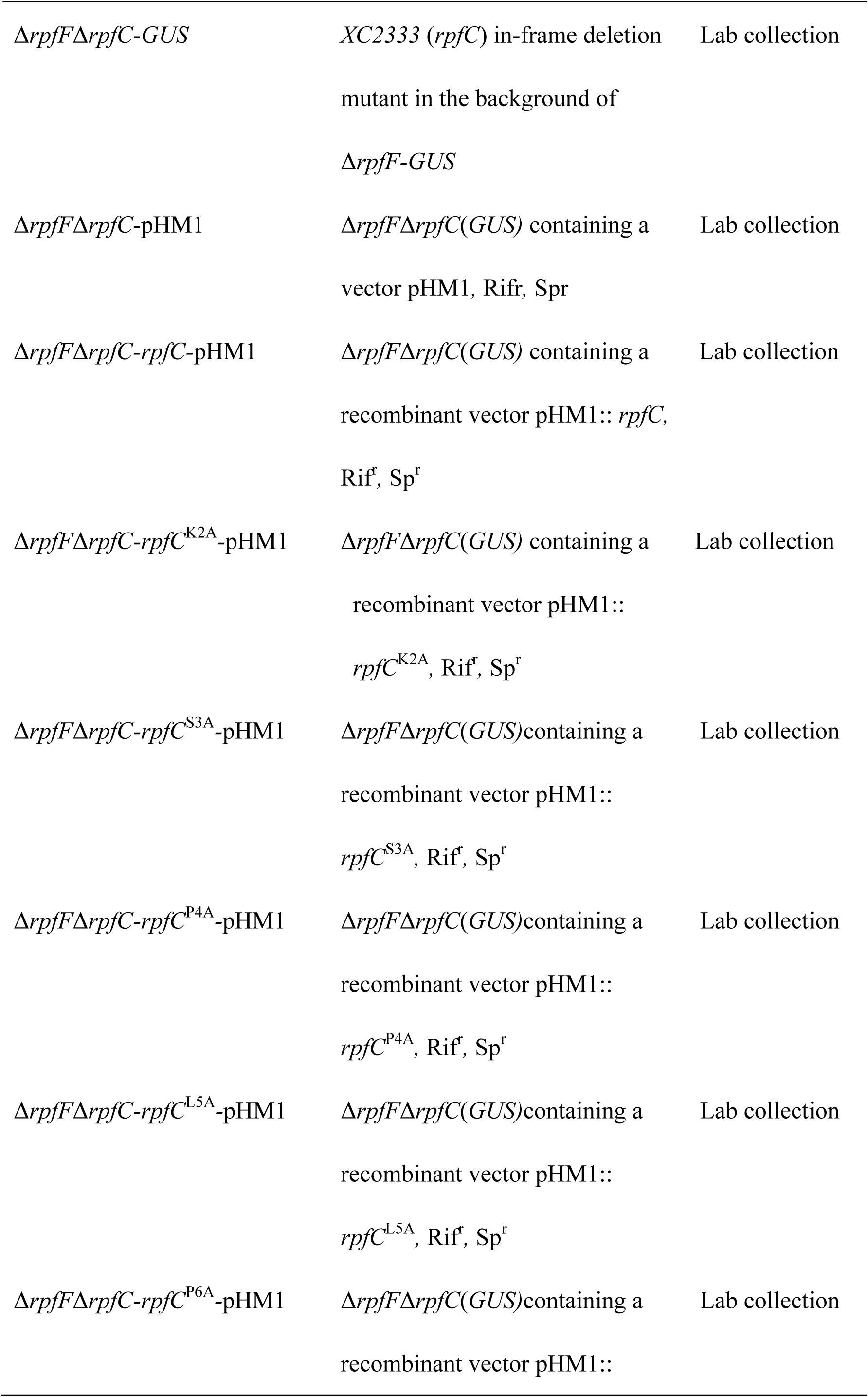
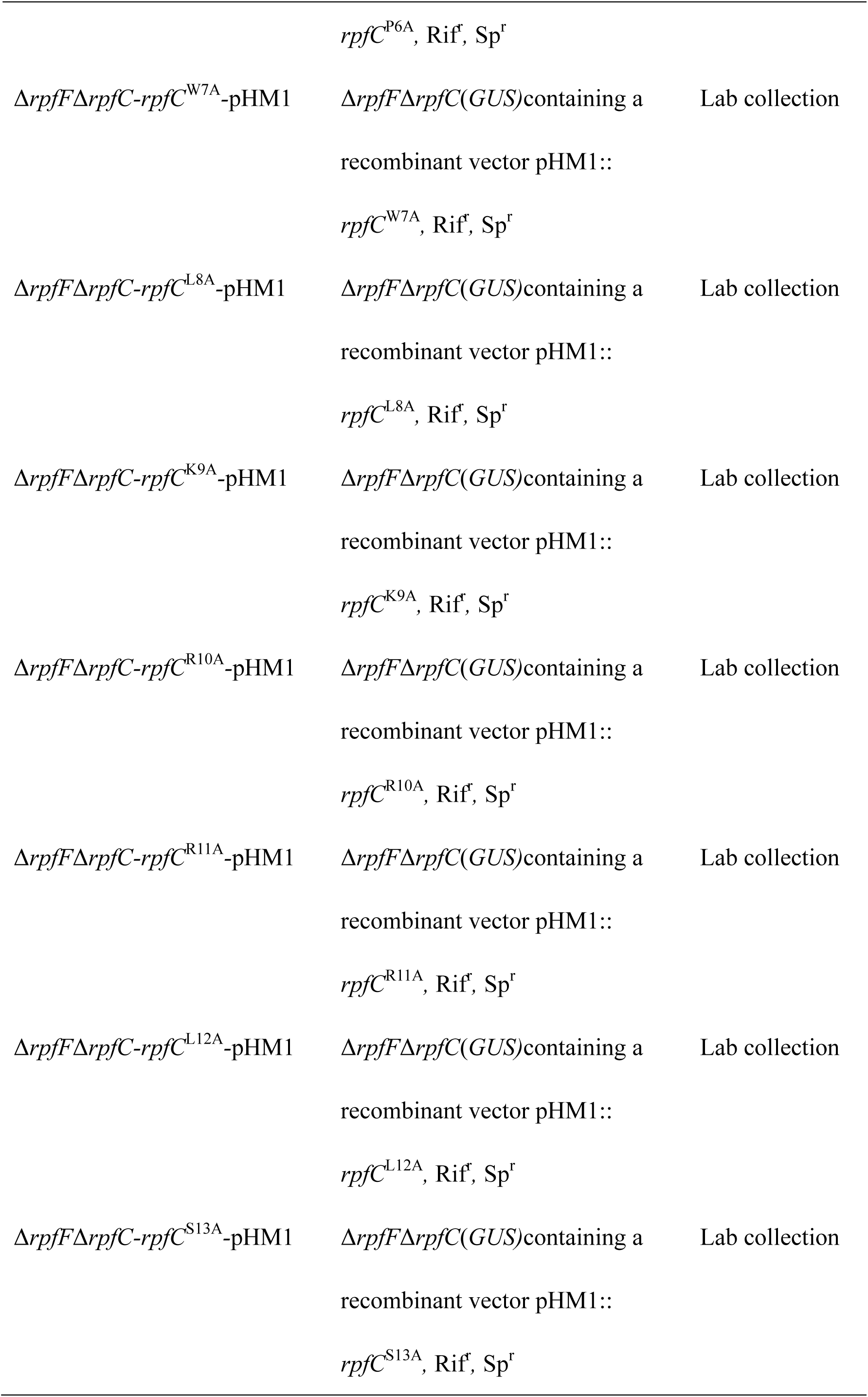
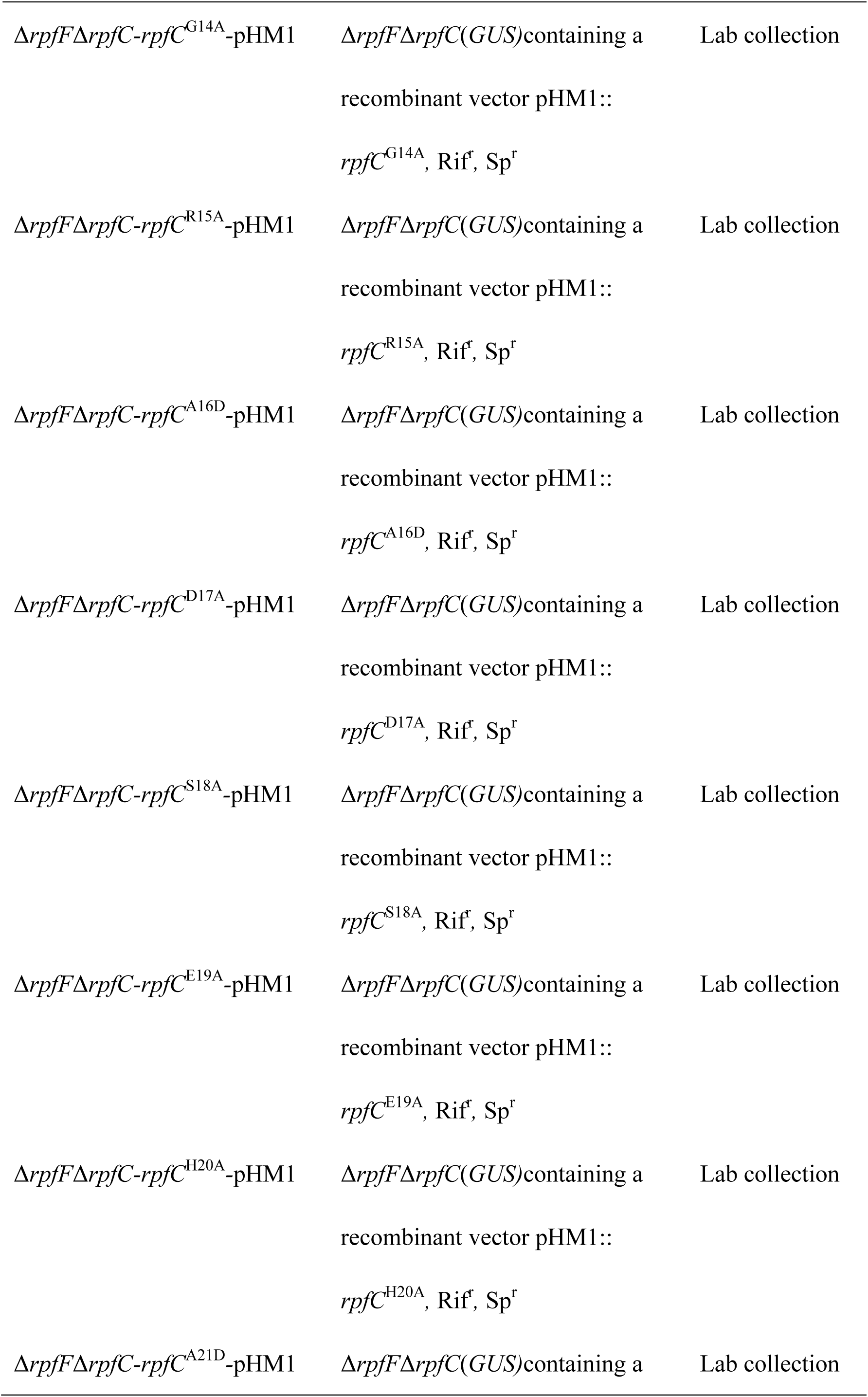
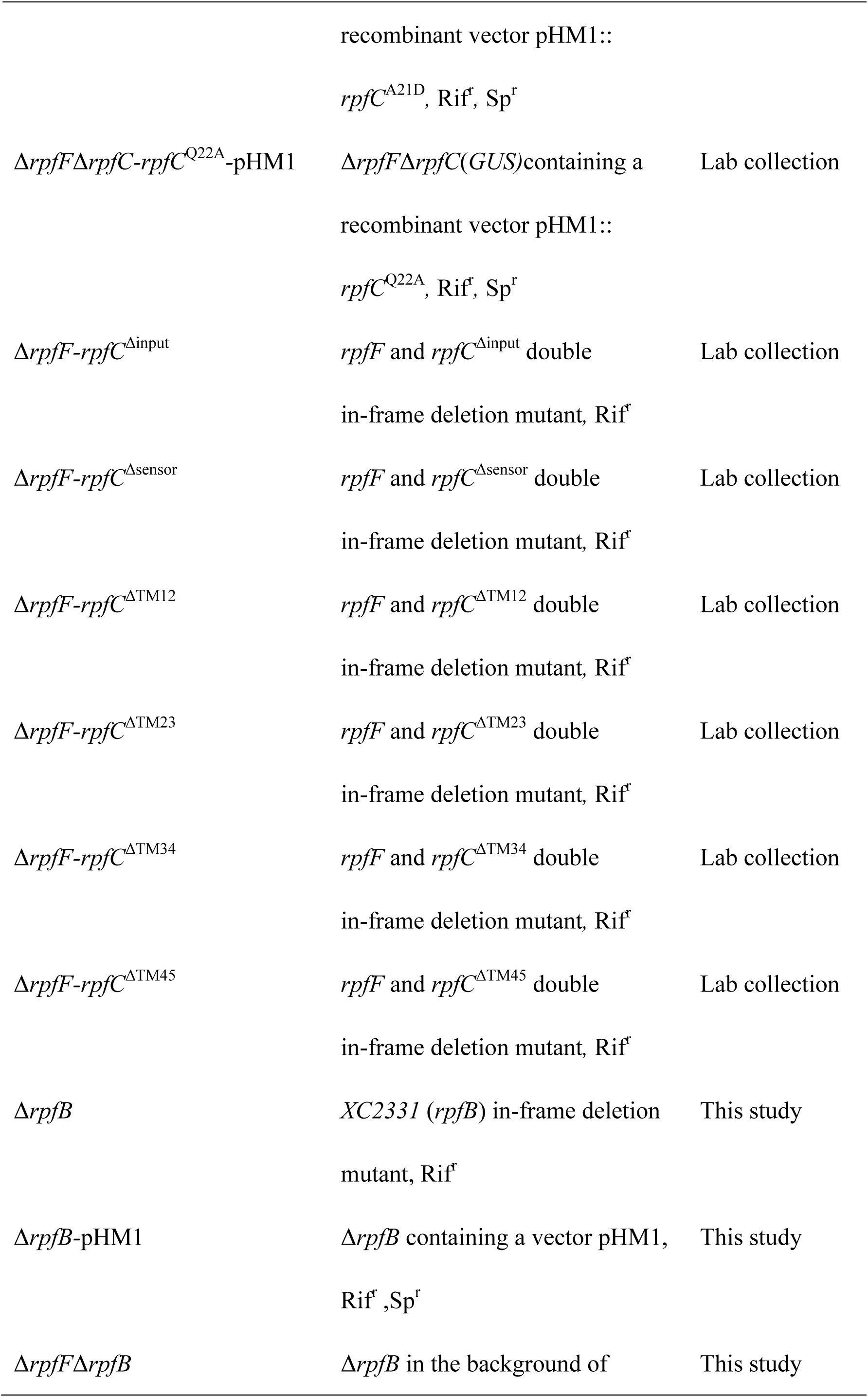
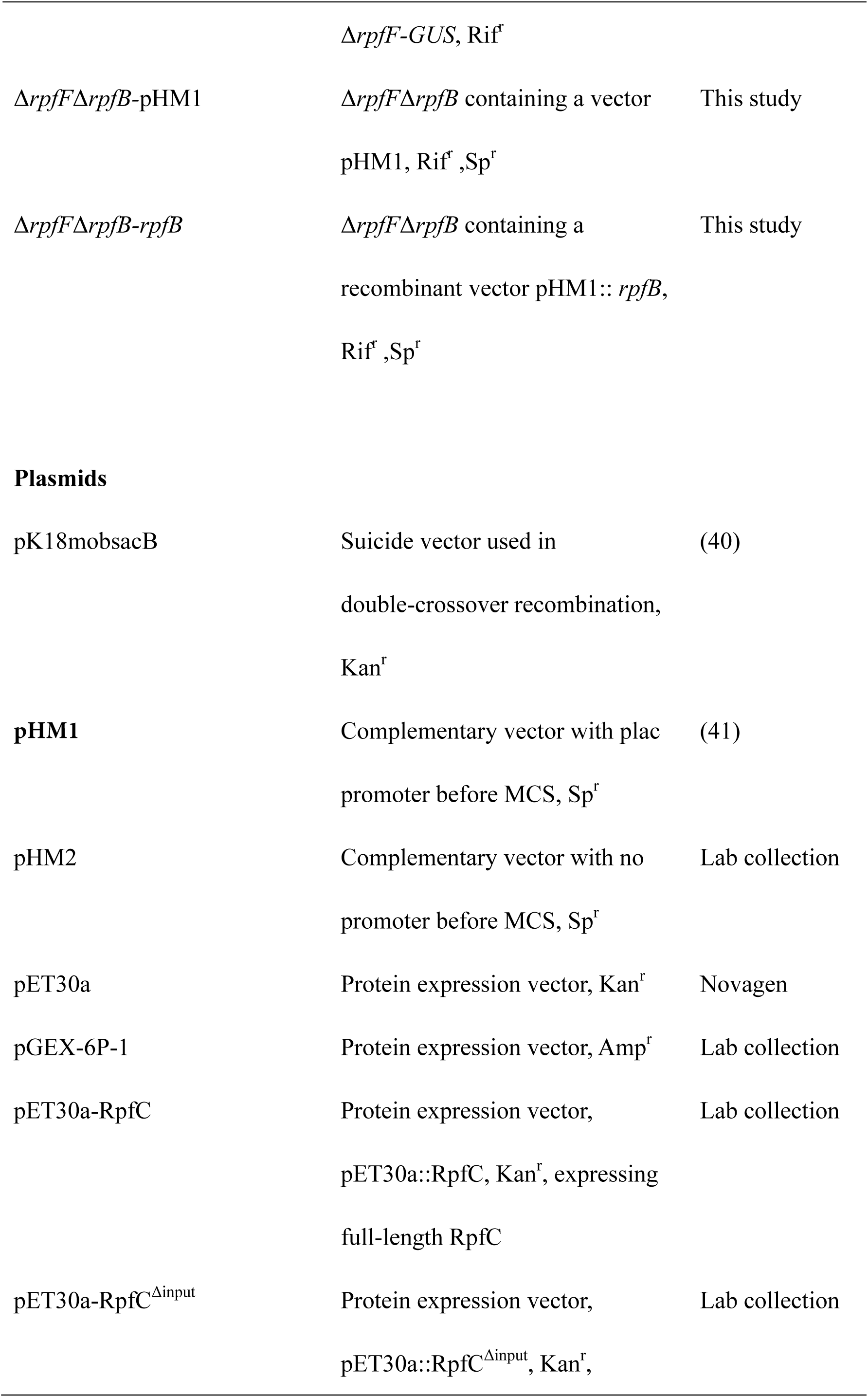
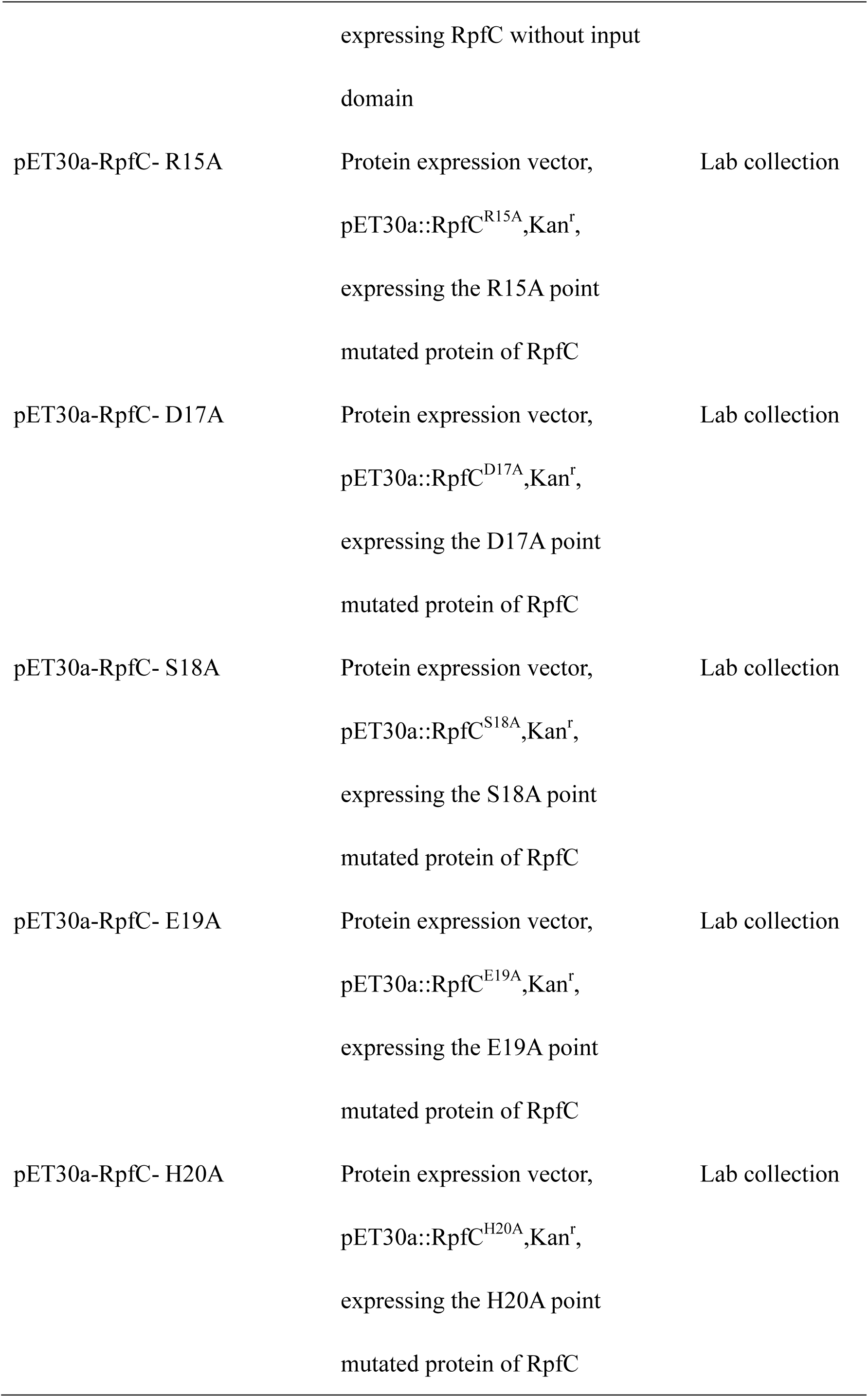
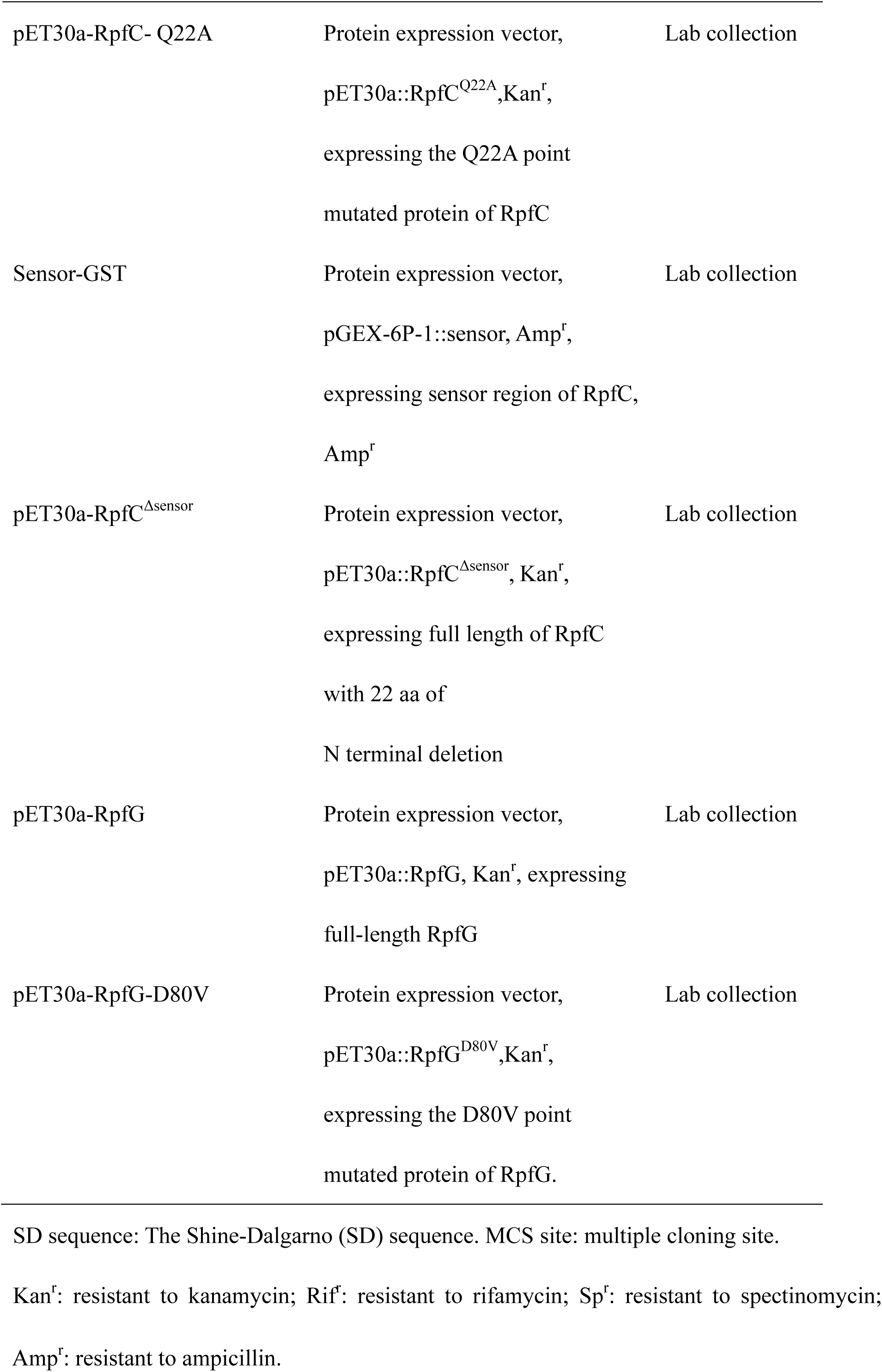
Bacterial strains and plasmids used in this study

| Strain or plasmid | Genotype or description | Source |
| --- | --- | --- |
| **Bacterial strains** |  |  |
| *Escherichia coli* strains |  |  |
| *E. coli* DH5α | *fhuA2* $\Delta$(*argF-lacZ*)U169 *phoA*<br>*glnV44* Φ80 $\Delta$(*lacZ*)M15 *gyrA96*<br>*recA1 relA1 endA1 thi-1 hsdR17* | Lab collection |
| **E. coli** BL21(DE3) | **E. coli B F<sup>-</sup> dcm omp<sup>T</sup> hsd(rB<sup>-</sup> mB<sup>-</sup>)gal $\lambda$ (DE3)** | Lab collection |
| *X. campestris* pv. *campestris* |  |  |
| *Xcc* 8004 | **Wild type strain (WT), Rif<sup>r</sup>** | Lab collection |
| Δ*rpfF* | *XC2332* (*rpfF*) in-frame deletion mutant, Rif | Lab collection |
| Δ*rpfF*-*GUS*, | a *gusE* gene and its own SD sequence was fused into the DNA region between *PengXCA* promoter and *engXCA* in the background of Δ*rpfF* | Lab collection |
| Δ*rpfC* | *XC2333* (*rpfC*) in-frame deletion mutant, Rif | Lab collection |
| $\Delta rpfF \Delta rpfC$-GUS | *XC2333 (rpfC)* in-frame deletion<br>mutant in the background of<br>$\Delta rpfF$-GUS | Lab collection |
| $\Delta rpfF \Delta rpfC$-pHM1 | $\Delta rpfF \Delta rpfC$(GUS) containing a<br>vector pHM1, Rif<sup>r</sup>, Spr | Lab collection |
| $\Delta rpfF \Delta rpfC$-*rpfC*-pHM1 | $\Delta rpfF \Delta rpfC$(GUS) containing a<br>recombinant vector pHM1::*rpfC*,<br>Rif<sup>r</sup>, Sp<sup>r</sup> | Lab collection |
| $\Delta rpfF \Delta rpfC$-*rpfC*<sup>K2A</sup>-pHM1 | $\Delta rpfF \Delta rpfC$(GUS) containing a<br>recombinant vector pHM1::<br>*rpfC*<sup>K2A</sup>, Rif<sup>r</sup>, Sp<sup>r</sup> | Lab collection |
| $\Delta rpfF \Delta rpfC$-*rpfC*<sup>S3A</sup>-pHM1 | $\Delta rpfF \Delta rpfC$(GUS)containing a<br>recombinant vector pHM1::<br>*rpfC*<sup>S3A</sup>, Rif<sup>r</sup>, Sp<sup>r</sup> | Lab collection |
| $\Delta rpfF \Delta rpfC$-*rpfC*<sup>P4A</sup>-pHM1 | $\Delta rpfF \Delta rpfC$(GUS)containing a<br>recombinant vector pHM1::<br>*rpfC*<sup>P4A</sup>, Rif<sup>r</sup>, Sp<sup>r</sup> | Lab collection |
| $\Delta rpfF \Delta rpfC$-*rpfC*<sup>L5A</sup>-pHM1 | $\Delta rpfF \Delta rpfC$(GUS)containing a<br>recombinant vector pHM1::<br>*rpfC*<sup>L5A</sup>, Rif<sup>r</sup>, Sp<sup>r</sup> | Lab collection |
| $\Delta rpfF \Delta rpfC$-*rpfC*<sup>P6A</sup>-pHM1 | $\Delta rpfF \Delta rpfC$(GUS)containing a<br>recombinant vector pHM1:: | Lab collection |
|  | *rpfC<sup>P6A</sup>*, Rif<sup>r</sup>, Sp<sup>r</sup> |  |
| $\Delta rpfF\Delta rpfC$-*rpfC<sup>W7A</sup>*-pHM1 | $\Delta rpfF\Delta rpfC$(*GUS*)containing a recombinant vector pHM1::<br>*rpfC<sup>W7A</sup>*, Rif<sup>r</sup>, Sp<sup>r</sup> | Lab collection |
| $\Delta rpfF\Delta rpfC$-*rpfC<sup>L8A</sup>*-pHM1 | $\Delta rpfF\Delta rpfC$(*GUS*)containing a recombinant vector pHM1::<br>*rpfC<sup>L8A</sup>*, Rif<sup>r</sup>, Sp<sup>r</sup> | Lab collection |
| $\Delta rpfF\Delta rpfC$-*rpfC<sup>K9A</sup>*-pHM1 | $\Delta rpfF\Delta rpfC$(*GUS*)containing a recombinant vector pHM1::<br>*rpfC<sup>K9A</sup>*, Rif<sup>r</sup>, Sp<sup>r</sup> | Lab collection |
| $\Delta rpfF\Delta rpfC$-*rpfC<sup>R10A</sup>*-pHM1 | $\Delta rpfF\Delta rpfC$(*GUS*)containing a recombinant vector pHM1::<br>*rpfC<sup>R10A</sup>*, Rif<sup>r</sup>, Sp<sup>r</sup> | Lab collection |
| $\Delta rpfF\Delta rpfC$-*rpfC<sup>R11A</sup>*-pHM1 | $\Delta rpfF\Delta rpfC$(*GUS*)containing a recombinant vector pHM1::<br>*rpfC<sup>R11A</sup>*, Rif<sup>r</sup>, Sp<sup>r</sup> | Lab collection |
| $\Delta rpfF\Delta rpfC$-*rpfC<sup>L12A</sup>*-pHM1 | $\Delta rpfF\Delta rpfC$(*GUS*)containing a recombinant vector pHM1::<br>*rpfC<sup>L12A</sup>*, Rif<sup>r</sup>, Sp<sup>r</sup> | Lab collection |
| $\Delta rpfF\Delta rpfC$-*rpfC<sup>S13A</sup>*-pHM1 | $\Delta rpfF\Delta rpfC$(*GUS*)containing a recombinant vector pHM1::<br>*rpfC<sup>S13A</sup>*, Rif<sup>r</sup>, Sp<sup>r</sup> | Lab collection |
| $\Delta rpfF\Delta rpfC$-$rpfC^{G14A}$-pHM1 | $\Delta rpfF\Delta rpfC(GUS)$containing a recombinant vector pHM1::<br>$rpfC^{G14A}$, Rif<sup>r</sup>, Sp<sup>r</sup> | Lab collection |
| $\Delta rpfF\Delta rpfC$-$rpfC^{R15A}$-pHM1 | $\Delta rpfF\Delta rpfC(GUS)$containing a recombinant vector pHM1::<br>$rpfC^{R15A}$, Rif<sup>r</sup>, Sp<sup>r</sup> | Lab collection |
| $\Delta rpfF\Delta rpfC$-$rpfC^{A16D}$-pHM1 | $\Delta rpfF\Delta rpfC(GUS)$containing a recombinant vector pHM1::<br>$rpfC^{A16D}$, Rif<sup>r</sup>, Sp<sup>r</sup> | Lab collection |
| $\Delta rpfF\Delta rpfC$-$rpfC^{D17A}$-pHM1 | $\Delta rpfF\Delta rpfC(GUS)$containing a recombinant vector pHM1::<br>$rpfC^{D17A}$, Rif<sup>r</sup>, Sp<sup>r</sup> | Lab collection |
| $\Delta rpfF\Delta rpfC$-$rpfC^{S18A}$-pHM1 | $\Delta rpfF\Delta rpfC(GUS)$containing a recombinant vector pHM1::<br>$rpfC^{S18A}$, Rif<sup>r</sup>, Sp<sup>r</sup> | Lab collection |
| $\Delta rpfF\Delta rpfC$-$rpfC^{E19A}$-pHM1 | $\Delta rpfF\Delta rpfC(GUS)$containing a recombinant vector pHM1::<br>$rpfC^{E19A}$, Rif<sup>r</sup>, Sp<sup>r</sup> | Lab collection |
| $\Delta rpfF\Delta rpfC$-$rpfC^{H20A}$-pHM1 | $\Delta rpfF\Delta rpfC(GUS)$containing a recombinant vector pHM1::<br>$rpfC^{H20A}$, Rif<sup>r</sup>, Sp<sup>r</sup> | Lab collection |
| $\Delta rpfF\Delta rpfC$-$rpfC^{A21D}$-pHM1 | $\Delta rpfF\Delta rpfC(GUS)$containing a | Lab collection |
|  | recombinant vector pHM1:: |  |
|  | *rpfC*<sup>A21D</sup>, Rif<sup>r</sup>, Sp<sup>r</sup> |  |
| $\Delta rpfF \Delta rpfC$-*rpfC*<sup>Q22A</sup>-pHM1 | $\Delta rpfF \Delta rpfC$(*GUS*)containing a | Lab collection |
|  | recombinant vector pHM1:: |  |
|  | *rpfC*<sup>Q22A</sup>, Rif<sup>r</sup>, Sp<sup>r</sup> |  |
| $\Delta rpfF$-*rpfC*<sup>Δinput</sup> | *rpfF* and *rpfC*<sup>Δinput</sup> double | Lab collection |
|  | in-frame deletion mutant, Rif<sup>r</sup> |  |
| $\Delta rpfF$-*rpfC*<sup>Δsensor</sup> | *rpfF* and *rpfC*<sup>Δsensor</sup> double | Lab collection |
|  | in-frame deletion mutant, Rif<sup>r</sup> |  |
| $\Delta rpfF$-*rpfC*<sup>ΔTM12</sup> | *rpfF* and *rpfC*<sup>ΔTM12</sup> double | Lab collection |
|  | in-frame deletion mutant, Rif<sup>r</sup> |  |
| $\Delta rpfF$-*rpfC*<sup>ΔTM23</sup> | *rpfF* and *rpfC*<sup>ΔTM23</sup> double | Lab collection |
|  | in-frame deletion mutant, Rif<sup>r</sup> |  |
| $\Delta rpfF$-*rpfC*<sup>ΔTM34</sup> | *rpfF* and *rpfC*<sup>ΔTM34</sup> double | Lab collection |
|  | in-frame deletion mutant, Rif<sup>r</sup> |  |
| $\Delta rpfF$-*rpfC*<sup>ΔTM45</sup> | *rpfF* and *rpfC*<sup>ΔTM45</sup> double | Lab collection |
|  | in-frame deletion mutant, Rif<sup>r</sup> |  |
| $\Delta rpfB$ | *XC2331* (*rpfB*) in-frame deletion | This study |
|  | mutant, Rif<sup>r</sup> |  |
| $\Delta rpfB$-pHM1 | $\Delta rpfB$ containing a vector pHM1, | This study |
|  | Rif<sup>r</sup>, Sp<sup>r</sup> |  |
| $\Delta rpfF \Delta rpfB$ | $\Delta rpfB$ in the background of | This study |
| | $\Delta rpfF$-GUS, Rif<sup>r</sup> | |
| $\Delta rpfF\Delta rpfB$-pHM1 | $\Delta rpfF\Delta rpfB$ containing a vector<br>pHM1, Rif<sup>r</sup>, Sp<sup>r</sup> | This study |
| $\Delta rpfF\Delta rpfB$-rpfB | $\Delta rpfF\Delta rpfB$ containing a<br>recombinant vector pHM1:: rpfB,<br>Rif<sup>r</sup>, Sp<sup>r</sup> | This study |
| **Plasmids** |  |  |
| pK18mobsacB | Suicide vector used in<br>double-crossover recombination,<br>Kan<sup>r</sup> | (40) |
| pHM1 | Complementary vector with plac<br>promoter before MCS, Sp<sup>r</sup> | (41) |
| pHM2 | Complementary vector with no<br>promoter before MCS, Sp<sup>r</sup> | Lab collection |
| pET30a | Protein expression vector, Kan<sup>r</sup> | Novagen |
| pGEX-6P-1 | Protein expression vector, Amp<sup>r</sup> | Lab collection |
| pET30a-RpfC | Protein expression vector,<br>pET30a::RpfC, Kan<sup>r</sup>, expressing<br>full-length RpfC | Lab collection |
| pET30a-RpfC<sup>Δinput</sup> | Protein expression vector,<br>pET30a::RpfC<sup>Δinput</sup>, Kan<sup>r</sup>, | Lab collection |
|  | expressing RpfC without input domain |  |
| pET30a-RpfC- R15A | Protein expression vector,<br>pET30a::RpfC<sup>R15A</sup>,Kan<sup>r</sup>,<br>expressing the R15A point mutated protein of RpfC | Lab collection |
| pET30a-RpfC- D17A | Protein expression vector,<br>pET30a::RpfC<sup>D17A</sup>,Kan<sup>r</sup>,<br>expressing the D17A point mutated protein of RpfC | Lab collection |
| pET30a-RpfC- S18A | Protein expression vector,<br>pET30a::RpfC<sup>S18A</sup>,Kan<sup>r</sup>,<br>expressing the S18A point mutated protein of RpfC | Lab collection |
| pET30a-RpfC- E19A | Protein expression vector,<br>pET30a::RpfC<sup>E19A</sup>,Kan<sup>r</sup>,<br>expressing the E19A point mutated protein of RpfC | Lab collection |
| pET30a-RpfC- H20A | Protein expression vector,<br>pET30a::RpfC<sup>H20A</sup>,Kan<sup>r</sup>,<br>expressing the H20A point mutated protein of RpfC | Lab collection |
| pET30a-RpfC- Q22A | Protein expression vector,<br>pET30a::RpfC<sup>Q22A</sup>, Kan<sup>r</sup>,<br>expressing the Q22A point<br>mutated protein of RpfC | Lab collection |
| Sensor-GST | Protein expression vector,<br>pGEX-6P-1::sensor, Amp<sup>r</sup>,<br>expressing sensor region of RpfC,<br>Amp<sup>r</sup> | Lab collection |
| pET30a-RpfC<sup>Δsensor</sup> | Protein expression vector,<br>pET30a::RpfC<sup>Δsensor</sup>, Kan<sup>r</sup>,<br>expressing full length of RpfC<br>with 22 aa of<br>N terminal deletion | Lab collection |
| pET30a-RpfG | Protein expression vector,<br>pET30a::RpfG, Kan<sup>r</sup>, expressing<br>full-length RpfG | Lab collection |
| pET30a-RpfG-D80V | Protein expression vector,<br>pET30a::RpfG<sup>D80V</sup>, Kan<sup>r</sup>,<br>expressing the D80V point<br>mutated protein of RpfG. | Lab collection |

4 SD sequence: The Shine-Dalgarno (SD) sequence. MCS site: multiple cloning site.
5 Kan<sup>r</sup>: resistant to kanamycin; Rif<sup>r</sup>: resistant to rifamycin; Sp<sup>r</sup>: resistant to spectinomycin;
6 $\text{Amp}^r$ : resistant to ampicillin.

**Table S2.**
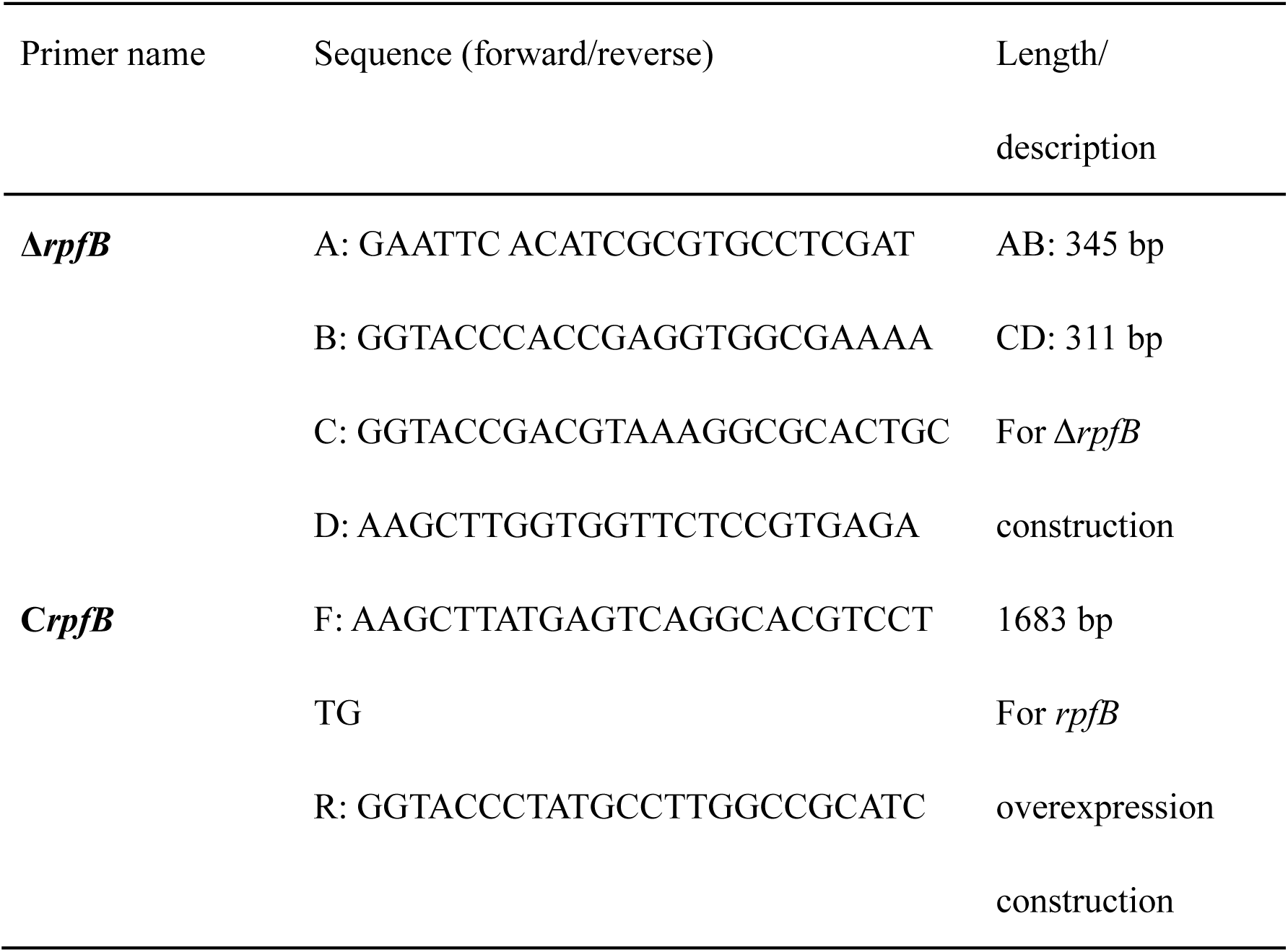
Primers used in this study

## Supplementary Figure Legends

**Figure S1.**
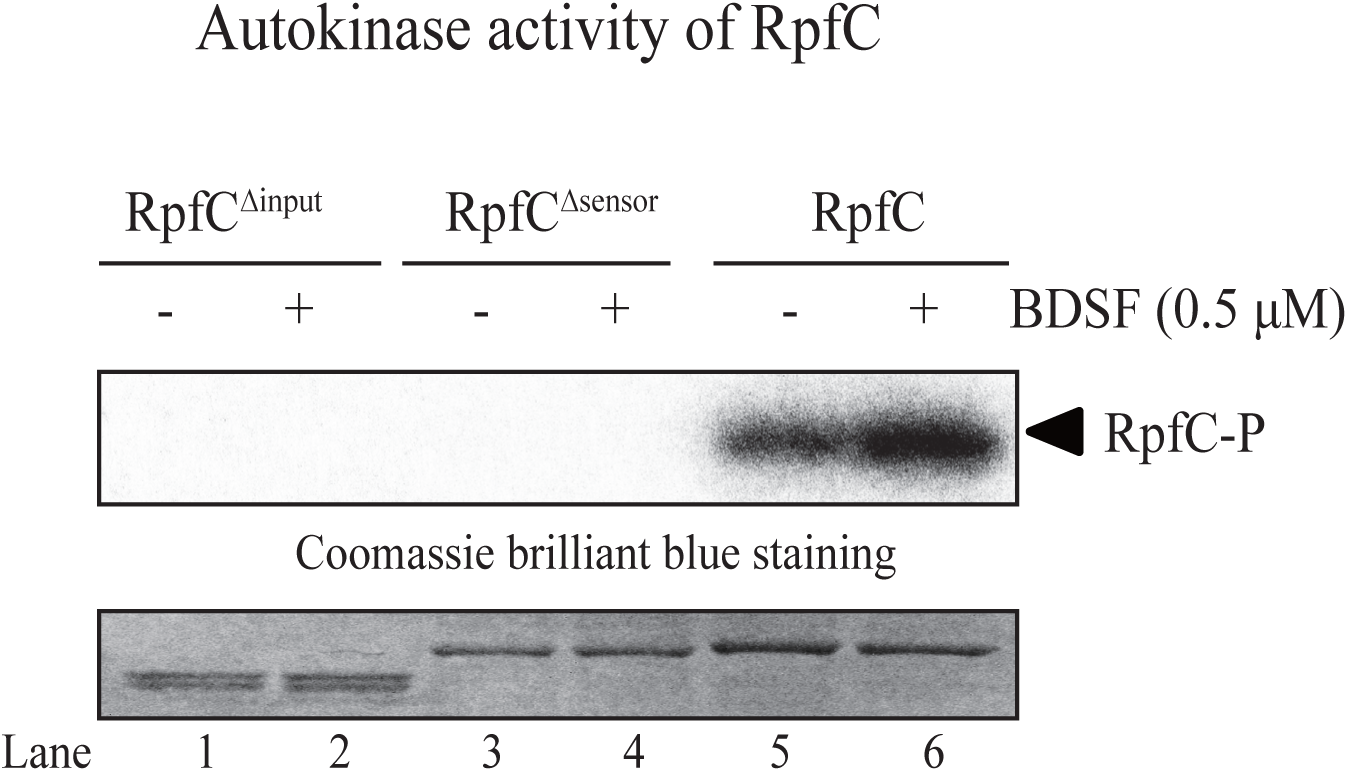
Full-length RpfC has autokinase activity. Autophosphorylation was detected using γ-[^32^P]ATP. The activities of the full-length RpfC embedded in liposomes as well as soluble, truncated RpfC without input or sensor regions were detected and recorded by autoradiography. Lower panels present RpfC stained with Coomassie brilliant blue, which served as loading controls. The experiment was repeated three times and representative results are presented.

**Figure S2.**
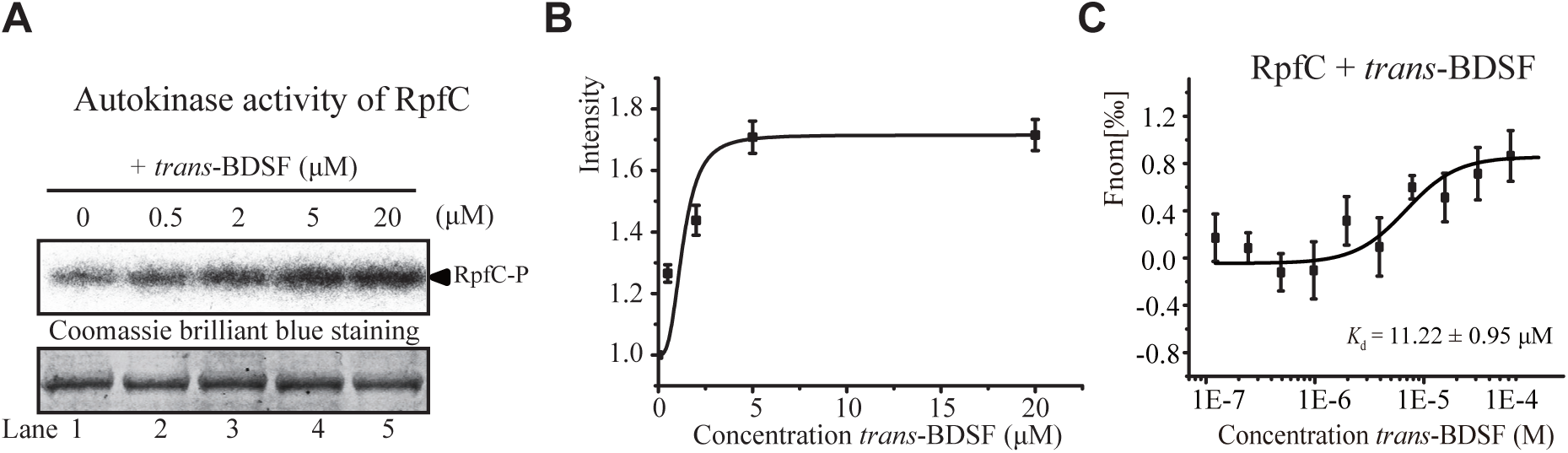
A high *trans*-BDSF concentration induces RpfC autokinase activity. (A) Autophosphorylation assay of RpfC. Various concentrations of *trans*-BDSF were added to the reaction mixture and the γ-[^32^P]ATP-labeled RpfC signal was detected and recorded by autoradiography. Lower panels present RpfC stained with Coomassie brilliant blue, which served as loading controls. (B) Dose-response analysis of RpfC autophosphorylation levels triggered by *trans*-BDSF. (C) Microscale thermophoresis assays revealed that *trans*-BDSF can bind to full-length RpfC liposomes, but at a low affinity. The *trans*-BDSF concentrations ranged from 25 to 2,000 μM. The solid curve presents the fit of the data points to the standard KD-Fit function. (A to C) Each experiment was repeated three times and black bars represent the standard deviation. *K*_d_ = dissociation constant.

**Figure S3.**
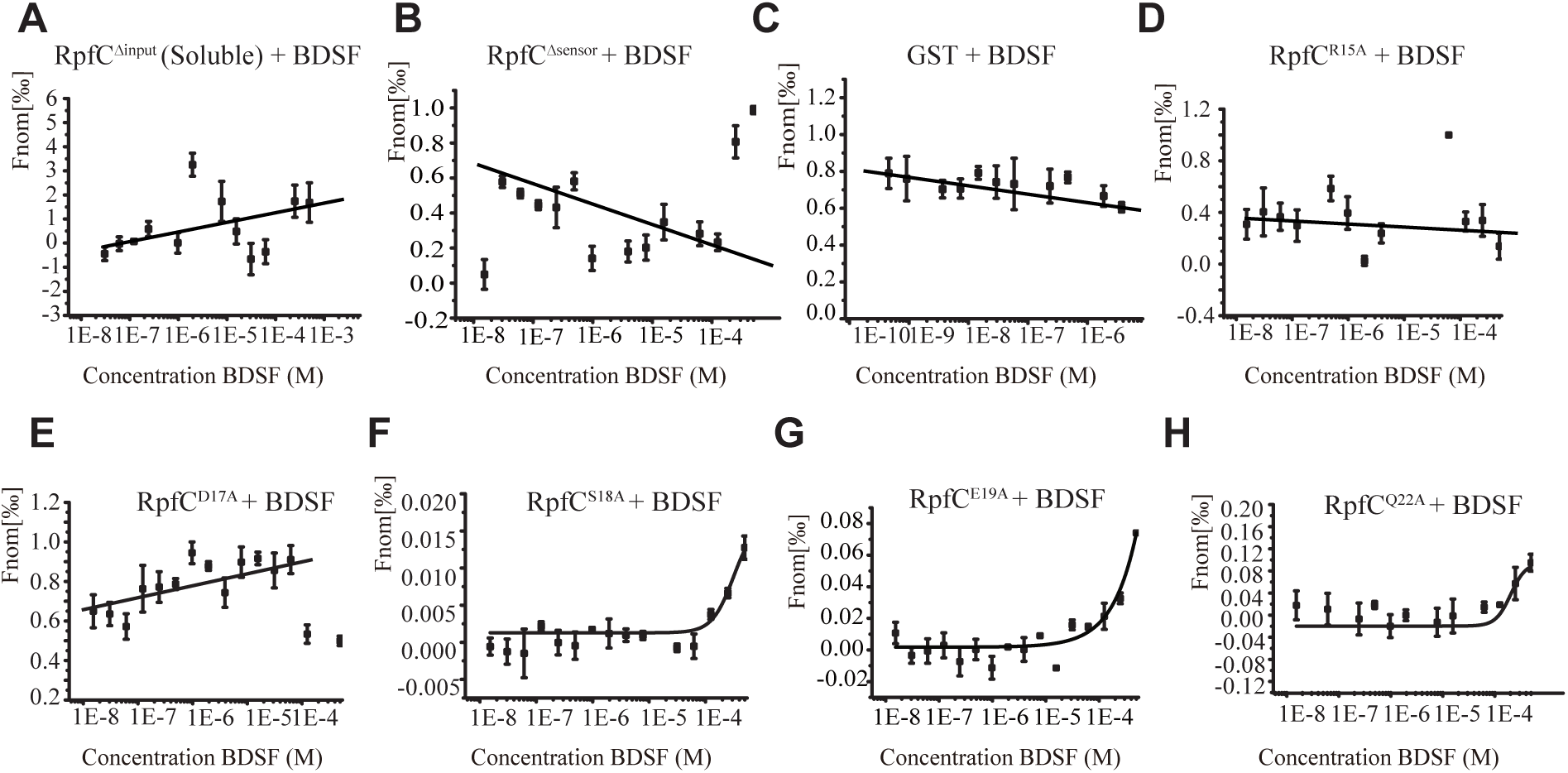
Substitution of essential residues in the RpfC sensor eliminates the binding between RpfC and BDSF. Microscale thermophoresis assays were performed to calculate the binding affinity. (A) BDSF did not bind to soluble RpfC lacking the input region. (B) BDSF did not bind to soluble RpfC lacking the sensor region. (C) BDSF did not bind to GST, which was used as a negative control for the purification of the sensor protein. (D to H) BDSF did not bind to recombinant RpfC with substituted amino acid residues in the sensor region: (D) RpfC^R15A^, (C) RpfC^D17A^, (F) RpfC^S18A^, (G) RpfC^E19A^, and (H) RpfC^Q22A^. The BDSF concentrations ranged from 25 to 2000 μM. The solid curve presents the fit of the data points to the standard KD-Fit function. Each experiment was repeated three times and black bars represent the standard deviation. *K*_d_ = dissociation constant.

